# A Systematic, Complexity-Reduction Approach to Dissect Microbiome: the Kombucha Tea Microbiome as an Example

**DOI:** 10.1101/2022.01.12.475982

**Authors:** Xiaoning Huang, Yongping Xin, Ting Lu

**Affiliations:** Department of Bioengineering, University of Illinois Urbana-Champaign, Urbana, IL, USA; Carl R. Woese Institute for Genomic Biology, University of Illinois Urbana-Champaign, Urbana, IL 61801, USA; College of Food Science and Nutritional Engineering, China Agricultural University, Beijing 100083, China; Department of Physics, University of Illinois Urbana-Champaign, Urbana, IL, USA; Center for Biophysics and Quantitative Biology, University of Illinois Urbana-Champaign, Urbana, IL 61801, USA; National Center for Supercomputing Applications, Urbana, IL 61801, USA

## Abstract

One defining goal of microbiome research is to uncover mechanistic causation that dictates the emergence of structural and functional traits of microbiomes. However, the extraordinary degree of ecosystem complexity has hampered the realization of the goal. Here we developed a systematic, complexity-reducing strategy to mechanistically elucidate the compositional and metabolic characteristics of microbiome by using the kombucha tea microbiome as an example. The strategy centered around a two-species core that was abstracted from but recapitulated the native counterpart. The core was convergent in its composition, coordinated on temporal metabolic patterns, and capable for pellicle formation. Controlled fermentations uncovered the drivers of these characteristics, which were also demonstrated translatable to provide insights into the properties of communities with increased complexity and altered conditions. This work unravels the pattern and process underlying the kombucha tea microbiome, providing a potential conceptual framework for mechanistic investigation of microbiome behaviors.

## Introduction

Microbiome populates the planet Earth, driving the growth of plants^1, 2^, biogeochemical cycling of elements^3, 4^, and health and disease of humans^5, 6^. Over the past decades, microbiome has gained explosive interest across disciplines from both academia and industry. To date, most efforts have focused on species cataloging^7^, composition-phenotype association^8, 9^ and microbiome-environment correlation^10^. These efforts yielded invaluable insights into ecosystem structure and function, reinforcing the need for microbiome research. Moving forward, an overarching goal is to dissect microbiome causation and mechanism^11, 12^. Specifically, required to be uncovered are the causes of specific microbiome traits and underlying mechanisms that drive the emergence of these traits. Tackling this challenge is important, because it will help to understand community structure and dynamics, predict the impacts of microbiome on habitats and design interventions for modulating ecosystem function^13, 14^.

To achieve the goal, one promising path is to dissect the metabolic underpinnings of members constituting a microbiome. Metabolism is a defining cellular process through which microbes acquire nutrient and energy; thus, its characteristics determine the growth of individual species. Through metabolism, cells also produce substances that are beneficial or deleterious to the growth of other species. Additionally, metabolism is often accompanied with the production of biomolecules that are bioactive to habitats (e.g., human, soil and plant). These molecules directly affect habitats, for instance, short-chain fatty acids produced by the gut microbiome shape the immune function and brain behavior of human^15–16^. Alternatively, they may remodel the physiochemical properties of the habitats, through which microbiome realizes indirect functional modulation. For example, extracellular polysaccharides secreted by probiotic bacteria trigger biofilm formation in the gastrointestinal tract, which promotes the host’ resistance to infection^17^. Thereby, targeting microbial metabolic underpinnings offers a systematic route to decode microbiome composition and function.

The pursuit of this path is, however, hindered by the intrinsic, remarkable complexity of native ecologies. For instance, the human gut microbiome consists of over 1,000 species and 100 trillion cells^18^; a teaspoon of healthy soil contains over 10,000 taxa members totaling up to 1 billion cells^19^. To circumvent the challenge, researchers have recently turned to microbiome cores^20–22^, simplified communities that are abstracted from native ecosystems but retaining their key structural and functional characteristics. Supporting the notion, studies have revealed a core gut microbiome across human population regardless of body weight^23^. Additionally, across soda lakes separated in distance, there is a collection of common microbes with similar structural patterns^24^. These simplified systems are approximations of native communities, providing a powerful alternative to study complex ecosystems.

Here we hypothesize to interrogate metabolic underpinnings of minimal cores as a causal and mechanistic strategy to elucidate microbiome structure and function. To test the hypothesis, we adopted the kombucha tea (KT) microbiome as our model ecosystem. Commonly called a symbiotic culture of bacteria and yeasts (SCOBY)^25^, the microbiome drives the fermentation of KT, a slightly sweet, acidic beverage with multiple health benefits^26, 27^. During the fermentation, the microbiome also produces floating pellicles at the air-liquid interface^28^. Compared to microbial ecologies in the soil and the human body, the KT microbiome is relatively simple in composition, easy to cultivate and amendable for quantification. Additionally, it involves species that are well characterized and feasible for perturbations. In fact, food microbiomes including those in kefir grain^29^, cheese rind^30–33^, wine^34^ and kimchi^35^ have been lately exploited as tractable platforms for studying community diversity, succession and niche partition^36^.

Our specific research started by characterizing the composition and metabolite patterns of the microbiome from commercially available KT drinks. We then used isolates to assemble 25 two-species consortia from which a minimal core was identified. Temporal fermentation showed that the core was convergent in its population composition, coordinated on temporal patterns of metabolites and capable of pellicle formation. Through comprehensive culturing of individual species under defined substrates, we obtained a casual and mechanistic understanding for the observed structural and functional traits of the core. We further showed that the knowledge from the core was translatable to account for the properties of communities with increased complexity and altered conditions. Together, our work illustrates the pattern and process underlying the composition and function of the KT microbiome, providing a promising conceptual framework for mechanistic investigation of microbiome behaviors.

## Results

### Characterization of the native KT microbiome

We set out to identify key structural and functional traits of the KT microbiome by performing fermentations with commercially available SCOBYs and black tea substrate supplemented with 50 g/L of sucrose (Methods). Each of the fermentations resulted in a light-brown broth and a floating, gel-like pellicle (Fig. 1a), which were analyzed in terms of their compositional diversity and metabolite abundance using amplicon sequencing and high-performance liquid chromatography respectively. Here, we considered metabolite profiles as a representation of microbiome function because chemical ingredients in KT broth are key factors conferring benefits^25, 27^.

**Figure 1.**
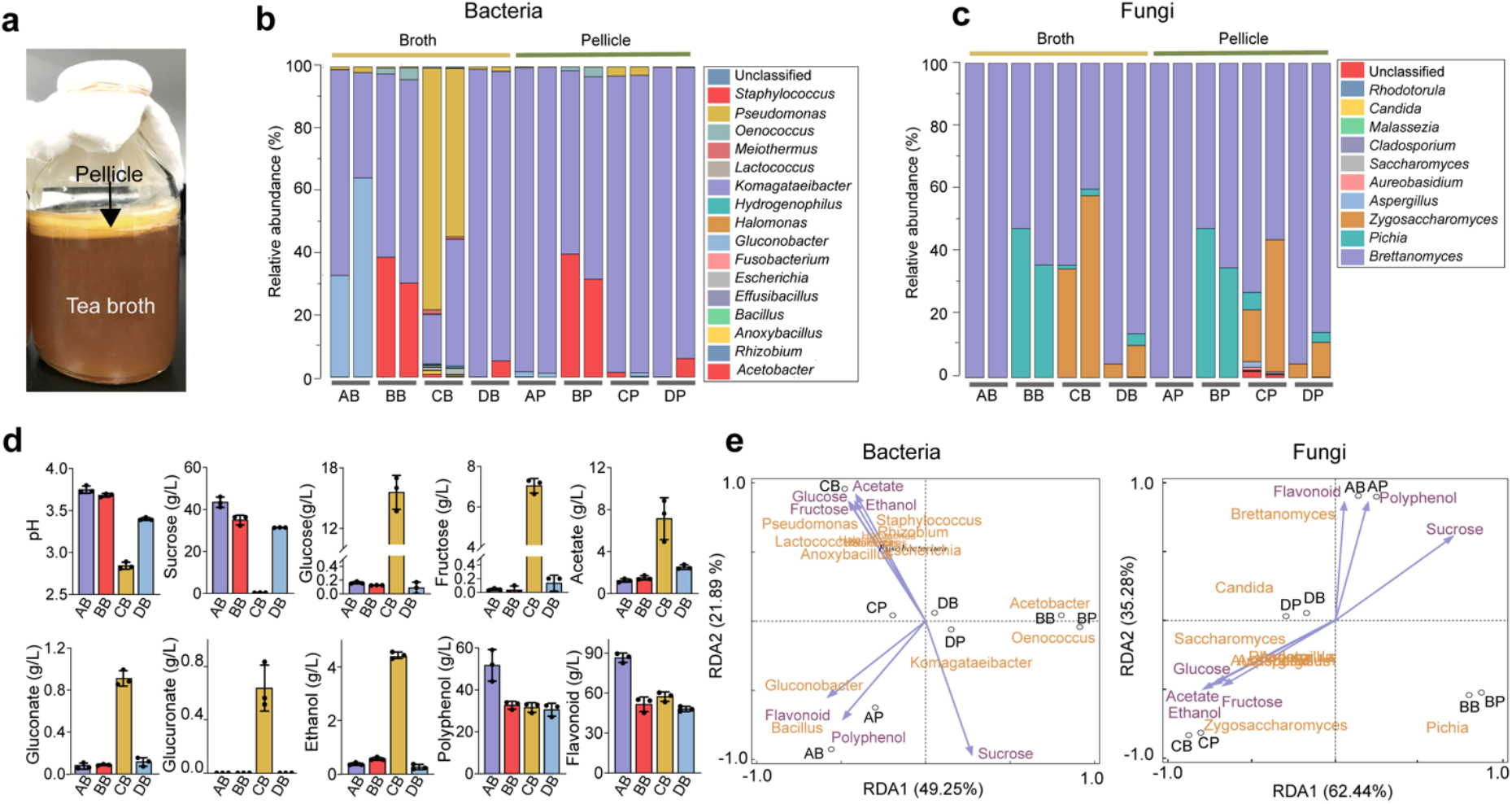
Characterization of the native KT microbiome. **a** Image of a typical kombucha tea fermentation containing both broth and pellicle. **b, c** Microbial composition in the broths and pellicles of different kombucha teas at the bacterial (**b**) and fungal (**c**) genus levels. For the four tea samples (A, B, C and D), their broths are named AB, BB, CB and DB whereas their pellicles are called AP, BP, CP and DP respectively. For each sample, two duplicates are presented. **d** Chemical properties of the kombucha tea broths. Measured variables include pH, sucrose, glucose, fructose, acetate, gluconate, glucuronate, ethanol, polyphenol and flavonoid. Bars and error bars correspond to means and s.d. **e** Correlation between microbial composition with biochemical substances in the KTs uncovered by redundancy analysis. Purple arrows represent metabolites, black circles represent different tea samples.

Our results showed that the microbiome had a relative low diversity, dominated by four bacterial genera, namely *Komagataeibacter*, *Acetobacter*, *Gluconobacter* and *Pseudomonas* (Fig. 1b), and three fungal genera including *Brettanomyces*, *Pichia* and *Zygosaccharomyces* (Fig. 1c). The bacteria and fungi also exhibited different context dependences: the composition of the former could vary significantly between the broth and pellicle of a single KT sample, such as samples A and C (AB (sample A’s broth) vs. AP (sample A’s pellicle), CB vs. CP) (Fig. 1b); by contrast, the composition of the latter remained consistent across broth and pellicle (Fig. 1c). Additionally, for bacteria, *Komagataeibacter* was the overall most predominant genus across samples and other genera were prevalent only in selected cases. For example, *Acetobacter* was prevalent in BB and BP, *Gluconobacter* was dominant in AB and *Pseudomonas* was predominant in CB. For fungi, *Brettanomyces* was predominant in all samples but, in samples B and C, *Pichia* and *Zygosaccharomyces* were also widespread. Thus, bacteria and fungi both served as constituting members of the microbiome, with *Komagataeibacter* and *Brettanomyces* being the dominant bacterial and fungal genus accordingly. This compositional pattern was consistent with previous reports although *Pseudomonas* was typically low in abundance^37–38^.

In parallel, we quantified the biochemical characteristics of KT broths, including pH, sugars, acids and tea-derived substances (Fig. 1d). The final pH values of samples A, B, D were around 3.6 while the pH of sample C was 2.8, all of which were in the reported range of a matured KT safe for human consumption^39^. The sucrose concentration dropped from 50 g/L to 30-40 g/L except for sample C whose sucrose was depleted. There were also trace amounts of glucose and fructose except for sample C containing a high level of the sugars. Acetate, gluconate and glucuronate were also detected, among which acetate had the highest concentration. Again, sample C was the outlier with a much higher level of acids. Since the concentration of gluconate was relatively low in our experiment and varied greatly across previous studies^26, 40^, we would not consider it as a characteristic metabolite. The fermentation also resulted in the accumulation of ethanol (∼0.5 g/L for samples A, B and D and 4.4 g/L for sample C). Two tea-derived compounds, polyphenol and flavonoid, were abundant (∼30 g/L and ∼50 g/L respectively). To reveal how these metabolites correlate with microbial composition, we performed redundancy analysis over the four samples (Fig. 1e).

From the above results, we drew three traits as the defining characteristics of the KT microbiome: first, it involves both bacteria and yeasts; second, it consumes sucrose with the synthesis of acetate, ethanol and a low level of glucose and fructose as the primary extracellular metabolites; third, it results in pellicle formation. These traits serve as the criteria for the identification of a proper microbiome core.

### Selection of a minimal core for the KT microbiome

To develop a correct core that recapitulates the native microbiome, we isolated a series of strains from the KT samples (Supplementary Tables 1,2). From the isolates, we selected 5 bacterial species, including *Komagataeibacter rhaeticus* (B_1_), *Komagataeibacter intermedius* (B_2_), *Gluconacetobacter europaeus* (B_3_), *Gluconobacter oxydans* (B_4_) and *Acetobacter senegalensis* (B_5_), and 5 fungal species, including *Brettanomyces bruxellensis* (Y_1_), *Zygosaccharomyces bailii* (Y_2_), *Candida sake* (Y_3_), *Lachancea fermentati* (Y_4_) and *Schizosaccharomyces pombe* (Y_5_), for synthesizing microbiome cores. Guided by the criterium that the KT microbiome contains both bacteria and fungi, we performed combinatorial mixing of the selected isolates, resulting in 25 two-species minimal core candidates with each involving one bacterial and one fungal species. To determine whether these candidates resemble the native, we conducted KT fermentation with these candidates and their corresponding 10 monocultures and, subsequently, quantified their microbial composition, extracellular metabolites and pellicle formation (Methods).

From colony forming units (CFU) counting (Fig. 2a), we found the bacterial and fungal species coexisted in all co-cultures as in the native KT microbiome. Additionally, in most cases, bacteria and yeasts had comparable relative abundances (<10 folds of difference) except for the combinations B_2_Y_3_ and B_4_Y_3_ whereby the bacteria were 100 times less than the yeast, suggesting these two combinations might not be the best candidates. For monocultures, the bacteria showed highly variable CFU while the yeasts yielded comparable CFU, indicating that bacteria varied greatly in sucrose utilization while yeasts were all stably capable.

**Figure 2.**
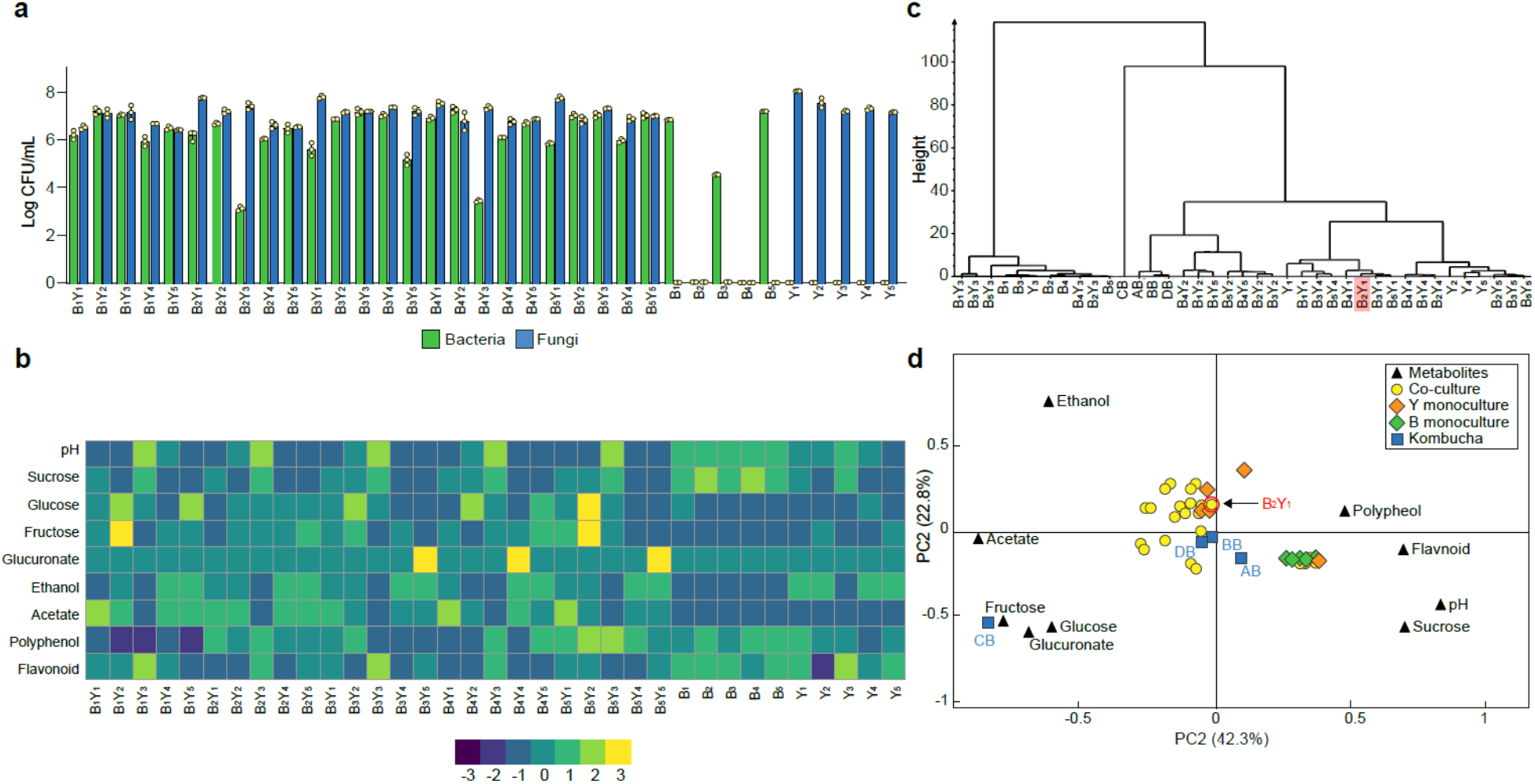
Population and metabolic quantification of two-species core candidates. **a** Colony forming units (CFU) counting of 25 two-species core candidates and 10 monoculture controls upon fermentation. Each core candidate is composed of one bacterial and one fungal species selected from the 10 isolates: B_1_ (*Komagataeibacter rhaeticus*), B_2_ (*Komagataeibacter intermedius*), B_3_ (*Gluconacetobacter europaeus*), B_4_ (*Gluconobacter oxydans*), B_5_ (*Acetobacter senegalensis*), Y_1_ (*Brettanomyces bruxellensis*), Y_2_ (*Zygosaccharomyces bailii*), Y_3_ (*Candida sake*), Y_4_ (*Lachancea fermentati*), and Y_5_ (*Schizosaccharomyces pombe*). Each monoculture control is one of the ten isolates. **b** Chemical property analysis of the core candidates and their controls. Heatmap is scaled by the values for each row. Measured variables include pH, sucrose, glucose, fructose, glucuronate, ethanol, acetate, polyphenol and flavonoid. **c** Hierarchical cluster analysis of the metabolic properties of the samples. The candidate B_2_Y_1_ is highlighted. **d** Principal component analysis of the metabolic properties. The candidate B_2_Y_1_ is circled in red.

By measuring pH, sugars, acids and tea-derived substances in the broths, we also obtained the biochemical characteristics of the candidates (Fig. 2b and Supplementary Table 3). The results showed that the co-cultures had comparable pH (∼ 3.5) except for the five involving Y_3_. The Y_3_-involving candidates also yielded a significantly higher level of residual sucrose and a significantly lower level of acetate and ethanol compared to others, suggesting that these candidates were unsuitable to serve as cores. The metabolite profiles of the monocultures showed that the yeasts alone could be sufficient for sucrose consumption. It also showed that acetate was produced primarily through co-cultures but not monocultures. To systematically evaluate the candidates, we performed hierarchical cluster analysis and principal component analysis over the metabolites to determine the similarities among the candidates and the four native samples (AB, BB, CB and DB). The hierarchical cluster analysis yielded three groups, one involving bacteria monocultures and Y_3_-involved mono- and co-cultures, another containing CB only, and the third including the rest (Fig. 2c). The principal component analysis showed that the co-cultures were all relatively close to the native microbiomes except for CB (Fig. 2d).

We further evaluated the candidates in terms of pellicle formation, the third characteristic of the native microbiome. The results showed that five co-cultures, B_2_Y_1_, B_2_Y_2_, B_2_Y_3_, B_2_Y_4_ and B_2_Y_5_, successfully produced pellicles during sucrose fermentation (data not shown).

Combining all three aspects of consideration, we chose B_2_Y_1_ as our minimal core of the KT microbiome for systematic, mechanistic investigation. Notably, Y_1_ (*B. bruxellensis*) was also the most predominant yeast species in the native samples (Fig. 1c and Supplementary Table 2).

### Compositional and metabolic dynamics of the core

To reveal the detailed traits of the selected core (B_2_Y_1_), we performed a set of fermentation experiments with different initial ratios (100:1, 10:1, 1:1, 1:10, and 1:100) while maintaining a constant total inoculation (2*10^6^ CFU/mL) (Methods). For all initial conditions, we found the bacterium B_2_ decreased in day 1 but increased afterwards with a declining magnitude of the growth rate (Fig. 3a and Supplementary Fig. 1a). By contrast, the yeast Y_1_ monotonically grew up with its rate reducing to null over time (Fig. 3b, Supplementary Fig. 1b). The population ratio of the two species showed that the community composition converged throughout the course of fermentation despite the variation of its initial ratio (Fig. 3c).

**Figure 3.**
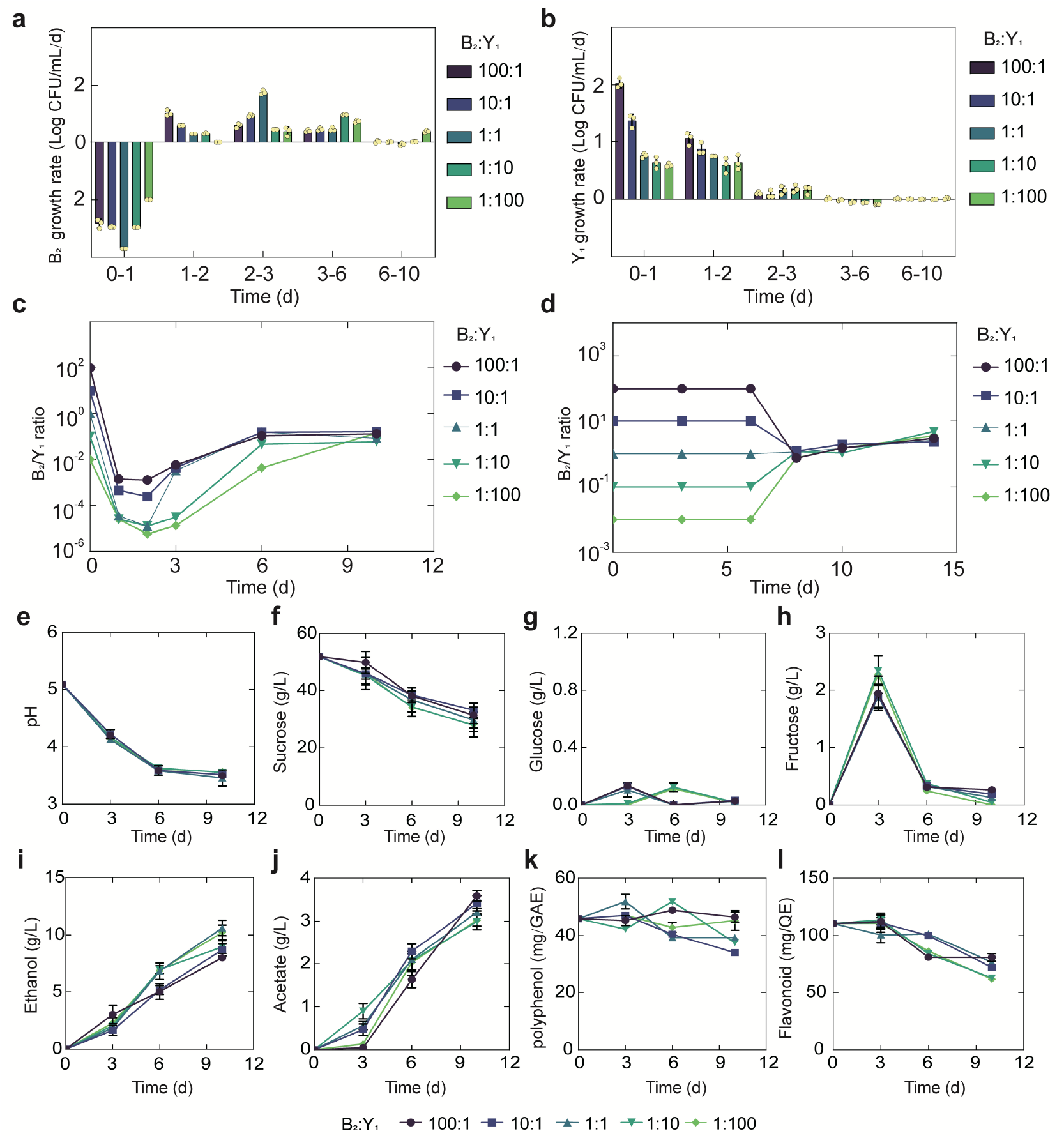
Temporal compositional and metabolic dynamics of the minimal core (B_2_Y_1_). **a**, **b** Growth rates of B_2_ (**a**) and Y_1_ (**b**) in tea broth during the KT fermentation with 50 g/L sucrose. **c** Bacterium-to-yeast population ratio of the microbes in broth. **d** Ratio of microbial populations in pellicle during the fermentation. **e-i** pH, carbon sources and metabolites during the fermentation driven by the core. Bars and error bars correspond to means and s.d.

The fermentation was also accompanied with the formation of pellicles (Supplementary Fig. 1c), which became visible after day 6 and grew continuously afterwards. Our CFU counting showed that, once pellicle formed, B_2_ and Y_1_ population densities remained relatively stable in the pellicles regardless of their initial abundance (Supplementary Fig. 1d,e). Meanwhile, their ratio converged to a fixed value (Fig. 3d) although the dry weight of the pellicles increased over time (Supplementary Fig. 1f). The convergence of composition in both broth and pellicle suggested that there were underlying forces that drove and stabilized community population dynamics.

Additionally, we quantified the temporal biochemical characteristics of the KT broth. Strikingly, although initial population ratios were varied across four orders of magnitude, each of the variables including pH, sugars, acids and tea-derived chemicals converged onto its own consensus pattern (Fig. 3e-l), akin to the convergence of composition in broth and pellicle. Specifically, regardless of the initial population composition, the pH dropped from 5.0 to 3.5 through fermentation (Fig. 3e), which was associated with continuous sucrose reduction (Fig. 3f). Throughout the process, glucose remained at a low level (∼0.1 g/L) (Fig. 3g) while fructose was relatively higher with a pulse-like profile (Fig. 3h). Acetate and ethanol on the other hand continued to accumulate during the fermentation (Fig. 3i,j). Polyphenol and flavonoids remained relatively stable with minor decrease (Fig. 3k,l). In the meanwhile, we found that throughout the fermentation process the temporal kinetics of different metabolites were coordinated. For example, continuous pH reduction (Fig. 3e) was in concert with sucrose drop (Fig. 3f), which was anti-correlated with the increase of ethanol (Fig. 3i) and acetate (Fig. 3j).

### Controlled fermentation assays yield causal claims for the core

To decode the mechanistic origins of the observed patterns, we investigated the metabolic processes of the constituting species (B_2_ and Y_1_) by conducting comprehensive monoculture fermentations with defined settings. Here, we focused on sucrose, glucose, fructose, ethanol and acetate as the primary biochemical substances of interest based on our measure of the KT broth and previous literature report^26, 27^. We used them alone and in combination as substrates to grow monocultures (Methods) and quantified the temporal profiles of key substances, pH, biomass growth and pellicle formation, resulting a total of 30 panels (Fig. 4).

**Figure 4.**
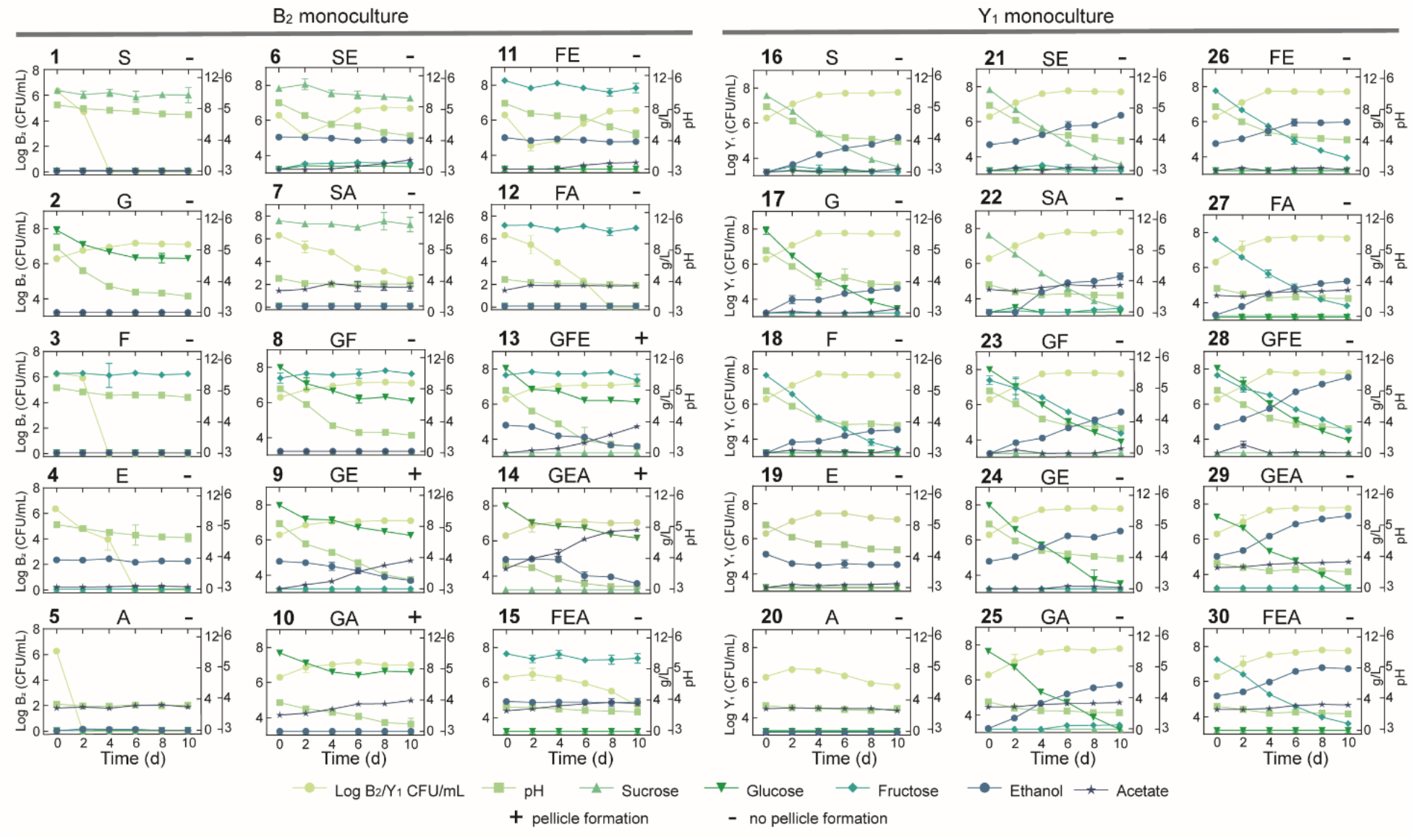
Comprehensive fermentation tests for the B_2_ and Y_1_ monocultures with different carbon sources. Sucrose (abbreviated as S, 10 g/L), glucose (G, 10 g/L), fructose (F, 10 g/L), ethanol (E, 50 mL/L) and acetate (A, 2 g/L) were used alone or in combination for fermentation. The number on top left of each panel is the label of the experiment. The letters on top middle of each panel indicate specific carbon sources used in the corresponding experiment. The + or – sign on the top right indicates whether a pellicle was formed during the fermentation. Bars and error bars correspond to means and s.d.

We harnessed the results of these panels to deduce biochemical conversion. As the starting carbon source, sucrose alone was not degradable by B_2_ as shown in panel 1 (abbreviated at P1) of Fig. 4 but consumable by Y_1_ with the production of a trace amount of glucose and fructose, ethanol accumulation, pH reduction and biomass growth (P16). Sucrose also showed weak hydrolysis in the presence of ethanol or acetate, which increased microbial survival (P6,7). Glucose and fructose were produced from sucrose hydrolysis primarily by Y_1_ (16) and minorly by ethanol and acetate (P6,7). Glucose was efficiently utilized by B_2_ for growth (P2) and by Y_1_ with biomass and ethanol accumulation (P17). Fructose was consumable for Y_1_ (P18), not B_2_ (P3), with ethanol and biomass production. Fructose was also slowly converted to glucose in the presence of ethanol, which supported B_2_ growth (P11). Ethanol was produced solely by Y_1_ during the metabolism of sucrose, glucose and fructose (P16,17,18), not by B_2_. Although ethanol alone was unusable by B_2_ (P4), it was consumed with glucose (P9), resulting in acetate production and pellicle formation without obvious growth benefits compared to glucose alone. It thus implied that ethanol was used an energy source for pellicle formation as previously reported^41^. Ethanol was also utilized by Y_1_ in a weak fashion to result in biomass and acetate production (P19). Acetate was produced primarily by B_2_ in the presence of multiple substrates (P6,9,13-15), particularly when glucose and ethanol were co-present (P9,13,14). In addition to B_2_, Y_1_ yielded a small amount of acetate with the consumption of sucrose, glucose, fructose or ethanol (P16-19). Acetate was additionally shown to minorly promote its own production by B_2_ (P10 vs. P2) and ethanol production by Y_1_ (P29,30 vs. P24,26).

Using the fermentation assays, we also inferred cellular tolerance to environmental stress. Comparison of the B_2_ and Y_1_ growth dynamics in single substrates showed that the yeast was more resistant than the bacterium to chemicals including ethanol and acetate (P4,5,19,20), which is another key factor that shapes community composition and metabolism. Additionally, the assays provided insight into pellicle formation. B_2_ monoculture was capable of pellicle production (P9,10,13,14) whereas Y_1_ was deficient under all conditions (P16-30). Moreover, comparison of the pellicle-forming conditions (P9,10,13,14) with single substrate conditions (P1-5) showed that efficient biofilm development required not only glucose but also ethanol or acetate as a co-substrate. Notably, although pellicle formation could occur in the presence of glucose as a sole carbon and energy source^42^ for certain species, at least for those we investigated, it required two substrates to produce pellicle.

The above findings were synthesized and subsequently integrated with reported metabolic reactions^43–46^ into a system-level diagram (Fig. 5), which involves major metabolic flows within each species, interspecies fluxes mediated by the environment, and regulatory effects from metabolites to fluxes. To illustrate its implications, we attempted to account for the observed compositional characteristics of the core. As the diagram showed, Y_1_ breaks down sucrose into glucose and fructose for its own growth, which also benefits B_2_ by sharing glucose. Additionally, Y_1_ secretes ethanol that is utilized by B_2_ when glucose is present. Thus, the core possesses a commensal relationship whereby Y_1_ provides two modes of benefits to B_2_. By design, such an interaction confers the stability and convergence of the ecosystem composition, thus providing a mechanistic driver for the population convergence in broth and pellicle (Fig. 3c,d). The results also elucidated three ways in which Y_1_ is more robust than B_2_: first, B_2_ relies on Y_1_ for glucose release; second, Y_1_ is more versatile for utilizing different substrates including sucrose, glucose, fructose and ethanol; third, Y_1_ has a higher tolerance to ethanol and acetate. These findings explained the temporal growth difference that B_2_ declined first before recovery while Y_1_ monotonically grew since the beginning of fermentation (Fig. 3a,b).

**Figure 5.**
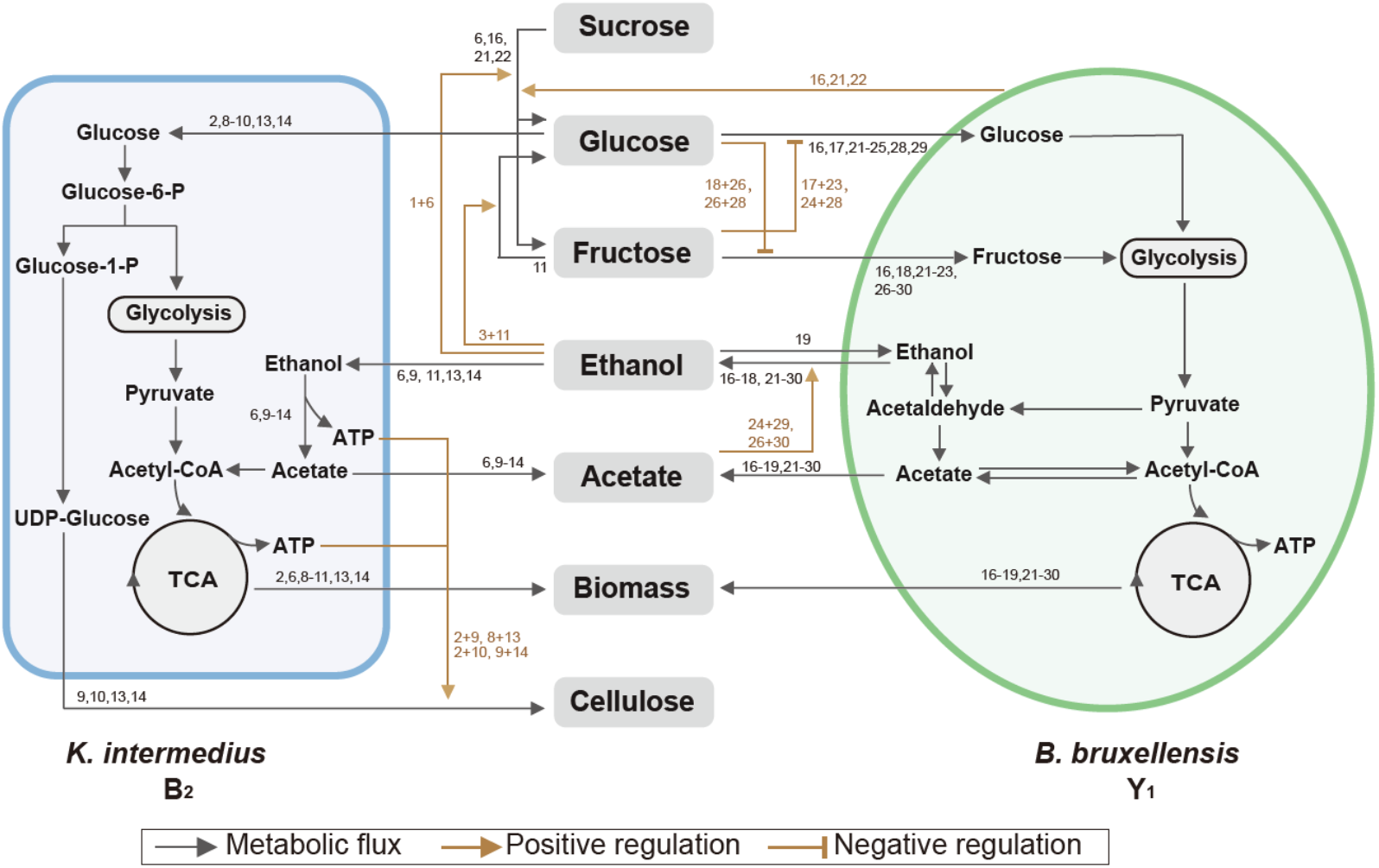
Summary of metabolic processes underlying the core. Black arrows refer to metabolic fluxes while brown arrows correspond to positive or negative regulatory interactions. The numbers associated with each arrow are the corresponding fermentation assays in Fig. 4 supporting the specific interaction.

Towards metabolic characteristics, the population of each species is a key determinant because total extracellular metabolites are determined by the productivity of individual cells multiplied by cell populations. Thus, for the same set of microbial species, the commensal interaction—which caused the convergence of population composition—also drove the convergence of temporal profiles of substrates, pH and metabolites as shown in Fig 3e-l. Additionally, although glucose and fructose were hydrolyzed simultaneously from sucrose, the former was consumed by both B_2_ and Y_1_ while the latter was useable exclusively for Y_1_, which resulted in a constant low level of glucose but a relatively higher level of fructose (Fig. 3g,h). Moreover, utilizations of glucose and fructose were accompanied with the release of ethanol and the both sugars were derived from sucrose hydrolysis; thus, ethanol increase was anti-correlated with sucrose decrease (Fig. 3f,i). Acetate was mainly produced by B_2_ in the presence of glucose and ethanol, both of which were converted directly or indirectly from sucrose; therefore, acetate accumulation was positively associated with sucrose consumption (Fig. 3f,j).

For pellicle formation, the diagram showed that B_2_ was solely responsible for pellicle formation. Meanwhile, it was Y_1_ that provided glucose and ethanol needed by B_2_. Such a cooperative relationship accounted for the findings that B_2_ or Y_1_ alone was deficient in pellicle formation and it needed the co-culture instead.

Relating to the bacterium-yeast symbiosis, some previous studies reported that the microbial social interactions are commensal while others concluded to be mutual^28, 47^. To resolve this debate, we conducted experiments to examine possible benefits from B_2_ to Y_1_. As certain yeasts were suggested to secret more invertase when co-cultured with cheaters^48^, we measured the invertase activity of Y_1_ in monoculture and in co-culture with B_2_ but did not find significant difference between the two conditions (*P*<0.05) (Supplementary Fig. 3). We also compared the growth and metabolites of Y_1_ in the B_2_Y_1_ co-culture and in monoculture with different initial ratios; however, the results showed that B_2_ did not affect either growth or metabolites except the increase of acetate which was produced by B_2_ (Supplementary Fig. 4). We additionally varied the B_2_ level while fixing Y_1_’s initial amount and altered Y_1_ while maintaining the initial B_2_. In both settings, Y_1_ growth was not affected by B_2_, and all metabolic variables, except acetate, exhibited the same patterns (Supplementary Figs. 5,6). Therefore, although we did not rule out the possibility of altered interactions across SCOBYs, our experiments demonstrated that, at least in our system, the symbiosis driving the community is commensal instead of mutual.

**Figure 6.**
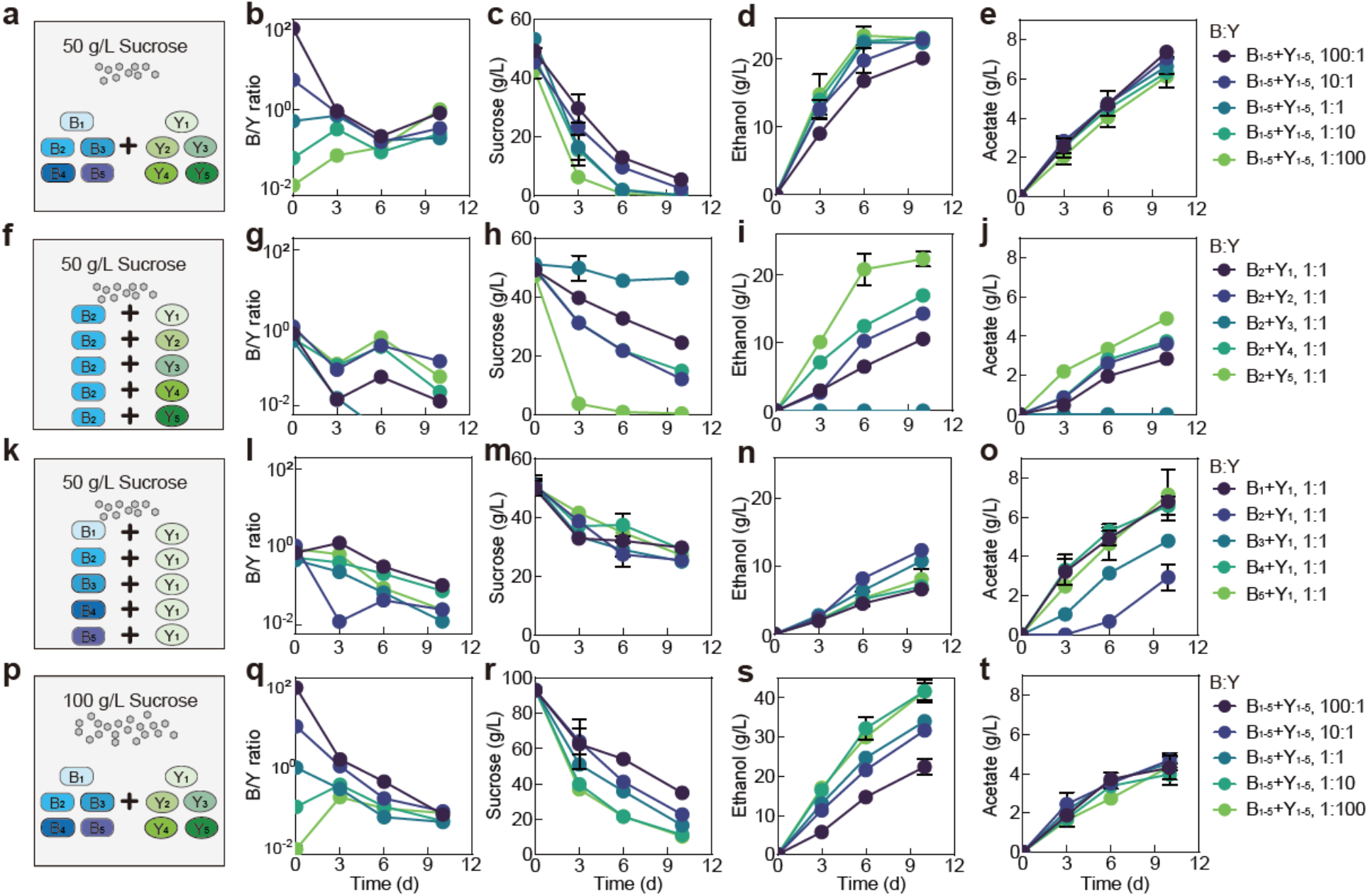
Fermentation by synthetic communities with increased complexity and altered conditions. **a,** Schematic illustration of a ten-species community involving B_1_-B_5_ and Y_1_-Y_5_ in a fermentation with 50 g/L of initial sucrose. **b-e** Population ratio (**b**), sucrose (**c**), ethanol (**d**) and acetate (**e**) throughout the course of the fermentation shown in **a**. **f** Schematic illustration of 5 two-species communities with each involving B_2_ and one of the yeasts (Y_1_-Y_5_) in a fermentation starting with 50 g/L sucrose. **g-j** Population ratio, sucrose, ethanol and acetate during the fermentation illustrated in **f**. **k** Schematic illustration of 5 two-species communities with each involving Y_1_ and one of the bacteria (B_1_-B_5_) in a fermentation with 50 g/L of initial sucrose. **i-o** Population ratio, sucrose, ethanol and acetate during the course of fermentation illustrated in **k**. **p** Schematic illustration of the ten-species community involving B_1_-B_5_ and Y_1_-Y_5_ in a fermentation starting with 100 g/L sucrose. **q-t** Population ratio, sucrose, ethanol and acetate during the fermentation depicted in **p**. Bars and error bars correspond to means and s.d.

### Insights into communities with increased complexity and varied conditions

We have thus far illustrated the casual claims for the two-species core, but do these findings provide implications for communities with different complexity and settings? To answer the question, we assembled a consortium of 10 species (B_1_, B_2_, B_3_, B_4_, B_5_, Y_1_, Y_2_, Y_3_, Y_4_ and Y_5_). Using the consortium, we performed fermentations with the same medium as previous (i.e., black tea substrate supplemented with 50 g/L sucrose) and using different initial total bacteria-to-yeast ratios while keeping all bacterial species even and all yeast species even (Fig. 6a).

In bulk, the ten-species community yielded the same patterns as the two-species core, including the overall compositional convergence compared to the initial structure (Fig. 6b), continued sucrose consumption (Fig. 6c), increase in ethanol and acetate (Fig. 6d,e), monotonic pH reduction (Supplementary Fig. 7b), consistent lowness of glucose (Supplementary Fig. 7c), pulse-like fructose profile (Supplementary Fig. 7d) and successful pellicle formation (data not shown). The similarity in patterns suggested that the core served as a good approximation of the ten-species consortium and that the knowledge from the simple core provided predictive insights into the behaviors of communities with an increased degree of complexity.

Meanwhile, in detail, the two ecosystems showed differences in specific profiles. In the ten-species community, the composition converged from day 0 to 6 but diverged at day 10 (Fig. 6b), different from the continuous convergence of the core (Fig. 3c,d). Compared to the core (Fig. 3), the ten-species community also yielded different metabolic patterns: its pH dropped faster (Supplementary Fig. 7b), sucrose was consumed quicker (Fig. 6c), fructose was more sensitive to initial conditions (Supplementary Fig. 7d), ethanol accumulated faster and nonlinearly with rapid production from day 0 to 6 followed by slower increase or crease from day 6 to 10 (Fig. 6d), and acetate increased faster (Fig. 6e).

We speculated that these differences arose from the variability of the metabolic capacities of members constituting the communities. To test the speculation, we repeated the fermentation with five two-species co-cultures involving B_2_ and different yeasts (Fig. 6f-j, Supplementary Fig. 7e-h). The results confirmed that the yeasts were highly variable in sucrose consumption with Y_5_ being the strongest and Y_3_ the weakest (Fig. 6h). Additionally, owing to the coordinated metabolism revealed via the core, rapid sucrose consumption by B_2_Y_5_ was associated with a relatively high B_2_ abundance, a large pulse of glucose and fructose, rapid ethanol and acetate production and quick pH drop (Fig. 6g-j, Supplementary Fig. 7f-h); by contrast, weak sucrose consumption by B_2_Y_3_ were accompanied with a low B_2_ abundance, an undetectable level of glucose and fructose, abolished ethanol and acetate production, and slow pH reduction (Fig. 6g- j, Supplementary Fig. 7f-h). We also performed fermentation using the co-cultures of Y_1_ with different bacterial species (Fig. 6k-o, Supplementary Fig. 7i-l). The five ecosystems showed a comparable sucrose consumption rate (Fig. 6m), suggesting that sucrose degradation was dictated primarily by yeast species although certain bacterial species (e.g. B_1_, B_3_, and B_5_ in Fig. 2a) could contribute. Meanwhile, the fermentations yielded varied ethanol and acetate patterns which were anti-correlated (Fig. 6n,o), suggesting that the ethanol-to-acetate conversion of the bacterial species were variable with B_1_ being the strongest and B_2_ the weakest.

From the above experiments, a mechanistic origin underlying the compositional and metabolic differences of the two- and ten-species communities emerged as following. Compared to the two-species core, the ten-species community had a higher overall sucrose consumption rate and a higher ethanol-to-acetate conversion rate which were averaged from the rates of the involved yeasts and bacteria. As a result, the ten-species community consumed sucrose faster (Fig. 6c), which subsequently led to a higher level of fructose, ethanol and acetate as well as a faster pH reduction (Fig. 6d,e, Supplementary Fig. 7b,d). Meanwhile, rapid sucrose degradation resulted in sucrose depletion in the middle of fermentation when the fermentation started with a high relative yeast abundance (e.g., 1:1, 1:10, 1:100) (Fig. 6c), which forced the yeast to metabolize fructose and ethanol instead of producing them. Under these scenarios, ethanol had a nonlinear pattern with rapid accumulation in the first few days and a slow increase or cease from day 6 to 10 (Fig. 6d). Meanwhile, as the commensal bacteria-yeast interaction relied primarily on the yeast to breakdown sucrose to provide glucose and ethanol for the bacteria, sucrose depletion also altered the strength of the symbiosis, which consequently shaped the dynamics of population convergence (Fig. 6b) because community dynamics was driven by the symbiosis.

Based on the finding that sucrose depletion shifted compositional and metabolic patterns, we hypothesized that, for the ten-species community, increasing sucrose availability could prevent sucrose depletion and hence drive its patterns closer to those of the core. We tested the hypothesis by performing the fermentation with 100 g/L sucrose (Fig. 6p-t, Supplementary Fig. 7m-p). Indeed, the results showed that the microbial composition continued to converge throughout the course of fermentation (Fig. 6q) instead of first convergence then divergence in Fig. 6b. Meanwhile, the continuous sucrose reduction (Fig. 6r) was accompanied with a faster and approximately linear ethanol increase (Fig. 6s), a lower rate of acetate accumulation (Fig. 6t), and a higher level of glucose and fructose (Supplementary Fig. 7o,p) compared to the 50 g/L sucrose case (Fig. 6b-d, Supplementary Fig. 7b-d). Notably, here the glucose and fructose patterns (Supplementary Fig. 7o,p) were still different from those of the core (Fig. 3g,h) because the ten-species community was much more efficient than the core for sucrose hydrolysis.

We further reasoned that the dependence of ecosystem characteristics on sucrose availability was not unique to the ten-species community and shall also apply to the two-species core. To test the reasoning, we performed the fermentation with the core using 5 g/L of sucrose (Supplementary Fig. 8). Remarkably, the composition converged in the first 3 days but diverged afterwards (Supplementary Fig. 8a) and ethanol started with linear increase initially but declined after day 3 (Supplementary Fig. 8h), similar to the case of the ten-species community with 50 g/L sucrose (Fig. 6b,d). Conversely, when we increased the initial sucrose concentration to 100 g/L (Supplementary Fig. 9), the compositional convergence of the core was restored (Supplementary Fig. 9a) and the ethanol profile became continuous accumulation (Supplementary Fig. 9h).

The results from the core and the ten-species community both informed that an increase in sucrose consumption results in a reduction or depletion in sucrose, an augmentation in the production of glucose, fructose, ethanol and acetate along with a reduction in environmental pH. Interestingly, such a relationship also explained the seemingly abnormal metabolite patterns of sample C, the outlier of the four native KT microbiome samples, that we observed at the beginning of our study (Fig. 1d). In that regard, our findings provided the mechanistic basis to understand the variations of metabolite patterns among the original microbiome samples.

Together, our experiments demonstrated that the knowledge from the minimal core offered predictive bulky insights into the traits of communities with varied system complexity and fermentation conditions. Meanwhile, the diversity in metabolic capacities, which was caused by the increase in species richness, accounted for the differences between the patterns of complex and minimal communities.

## Discussion

With rapid advances in species cataloging and correlation analysis, one remaining grand challenge in microbiome research is to uncover causal claims that dictate microbial composition and function^11–14, 49, 50^. In this work, we present the identification, characterization and utilization of a minimal core for elucidating the molecular mechanisms driving the KT microbiome. We showed that metabolic underpinnings specified the structural and metabolic characteristics of the core and also provided insights into the behaviors of communities with increased complexity and altered conditions.

Lying at the heart of our study is the reduction in system complexity, which involves three key steps: identification of a core simplified from a native microbiome but capable of resembling the native, characterization of metabolic underpinnings of the core, and extrapolation of the knowledge from the core to communities with altered complexity and conditions. Although this study focused exclusively on the KT microbiome, the strategy demonstrated here is not limited to the specific ecosystem. Given its systematic nature, we expect that it may be extendable to other microbial communities. In that regard, our strategy provides a promising solution to address system complexity, a major hurdle for mechanistic investigation of microbiome.

Notably, although minimal cores serve as attractive alternatives to complex ecosystems, they are not intended to substitute native microbiomes. With the increase of complexity, certain compositional and metabolic traits identified in a core may be altered in its native counterpart. Conversely, novel properties may emerge when species richness increases. Thus, minimal cores provide a point of entry to unlock the mechanistic behaviors of a community, which shall be followed by the analysis of the full system for systematic understanding. Meanwhile, defining a proper core is critical for successful implementation of our framework. In principle, a single microbiome can possess multiple cores depending on different selection criteria, such as abundance, temporal pattern and function. Recent studies suggested to utilize gene level analysis rather than organismal lineage for core identification^23, 51^. Nevertheless, future efforts in this direction are needed to fully realize the power of this community analysis strategy.

A major goal of the food industry is to improve food quality and flavor through the optimization of starter culture and fermentation process^52–54^. The synthetic core developed here successfully drove the KT fermentation, thereby serving as an effective, functional culture starter. Compared to the native microbiome, this well-defined, synthetic system offers a controllable platform to modulate the starter composition and metabolite secretion during fermentation. Therefore, the work also gives a potential solution to systematically tailor fermented foods with desired traits.

## Materials and Methods

### Kombucha tea fermentation

Black tea (Harney & Sons Fine Teas, Millerton, NY) was purchased as the tea substrate for the fermentation. The live starter culture SCOBY used as inoculum were obtained from 4 different commercial sources, referring to samples A, B, C and D. Kombucha tea was prepared as previously reported with minor modifications^55, 56^. Briefly, 1 L deionized water was boiled, added with 12 g/L black tea and allowed to infuse for 5 min. After removing the tea leaves, sucrose (50 g/L) was dissolved in hot tea. After cooling, the tea mixture was filtered through sterile sieve to 500 mL glass vessel with cotton and gauze caps. Then, 3.0% SCOBY and liquid broth (10% v/v) of the SCOBY samples were added to tea broth. The kombucha tea was incubated at 25°C for 14 days.

### Amplicon sequencing of 16S ribosomal RNA (rRNA) and ITSs

Pellicle samples were first treated with 200 mg/mL cellulase (Sigma-Aldrich, Milan, Italy) for 16 h, and sonicated in ice bath for 1 min using a probe sonicator (Model 505, Fisherbrand, USA) for 1 min. Then the samples were centrifuged at 6500 rpm at 4°C for 10 min. The cell pellets were used for DNA extraction. Total DNA extractions were performed for tea broth and pellicle samples using Quick-DNA Fecal/soil Microbe Miniprep kit (ZYMO Research Corp.) according to the manufacturer’s instructions.

16S rRNA gene and ITS amplicon sequencing library constructions and Illumina MiSeq sequencing were conducted by GENEWIZ, Inc. (South Plainfield, NJ, USA). Sequencing library was prepared using a MetaVx™ 16s rDNA Library Preparation kit and ITS-2 Library Preparation kit (GENEWIZ, Inc., South Plainfield, NJ, USA). Briefly, for each sample, 50 ng DNA was used to generate amplicons that cover the V3 and V4 hypervariable regions of bacteria and ITS-2 hypervariable region of fungi. Afterwards, each sample was added with indexed adapters. The barcoded amplicons were sequenced on the Illumina MiSeq platform using 2 × 250 paired-end (PE) configuration (Illumina, San Diego, CA, USA).

Raw sequence data was converted into FASTQ files and de-multiplexed using Illumina’s bcl2fastq 2.17 software. QIIME data analysis package was used for 16S rRNA and ITS rRNA data analysis^57^. All the reads (forward and reverse) were assigned to different samples based on barcode, and then truncated by cutting off the primer and barcode. After quality filtering^58^, the sequences were compared with the RDP Gold database to detect chimeric sequences using the UCHIME algorithm^59^. Subsequently, the effective sequences were grouped into operational taxonomic units (OTUs) using the clustering program VSEARCH (1.9.6) against the Silva 119 database for bacteria^60^ and the UNITE ITS database for fungi^61^, with pre-clustered at 97% of sequence identity. The Ribosomal Database Program (RDP) classifier was used to assign taxonomic category to all OTUs at a confidence threshold of 80%^62^.

### Species isolation and identification

For species isolation, kombucha tea broths were diluted and plated directly whereas pellicle samples were sonicated and digested before dilution as described above. Diluted samples were then inoculated in different selective media. For bacterial isolation, de Man, Rogosa and Sharpe medium (MRS), Mannitol medium^40^, and Glucose yeast extract calcium carbonate medium (GYC)^63^ were used in conjugation with 0.1% cycloheximide or 500 ug/mL natamycin for inhibiting fungi growth. Isolation of yeast species was carried out using the yeast extract peptone dextrose (YPD) medium supplemented with 100 mg/L chloramphenicol. Isolated species were identified by Sanger sequencing of the 16S and 26S rRNA gene regions, with the universal primers B-f (5′-AGAGTTTAGTCCTGGCTCAG-3′) and B-r (5′- AAGGAGGTGATCCAGCCGCA-3′) for bacteria^64^, and NL-1 (5΄- GCATATCAATAAGCGGAGGAAAAG-3΄) and NL-4 (5΄-GGTCCGTGTTTCAAGACGG-3΄) for yeasts^40^.

### Biochemical analyses

The pH was measured with a pH meter (AE150; Fisher Scientific, Waltham, MA) inserted directly into samples. Acetate, glucuronate and ethanol concentrations were determined by high performance liquid chromatography (HPLC, Agilent Technologies 1200 Series) equipped with a refractive index detector using a Rezex ROA Organic Acid H+ (8%) column (Phenomenex Inc. Germany). The column was eluted with 0.005 N of H_2_SO_4_ at a flow rate of 0.6 mL/min at 50°C^65^. Sucrose, glucose and fructose were analyzed using RCM Monosaccharide Ca^2+^ column (Phenomenex Inc., Germany). The column was eluted with deionized water at a flow rate of 0.6 mL/min at 80°C^66^. For gluconate detection, the gluconic acid Kit (Megazyme, Ireland) was used. The concentration of total polyphenols was measured by the Folin-Ciocalteu colorimetric method, with gallic acid as standard. The absorbance was measured at 765 nm and the results were expressed as mg of gallic acid equivalent (GAE) per mL of kombucha tea (mg GAE/mL)^67^. The total flavonoids were determined using an aluminum chloride assay using quercetin as standard. The absorbance was measured at 430 nm and the content was expressed as mg of quercetin equivalent (QE) per mL of kombucha tea (mg QE/mL)^68^. The invertase activity was determined according to the method described by Laurent *et al*^69^. The remaining sucrose was detected by HPLC as described above. The measurement of pellicle weight was based on the descriptions of Florea *et al.* using 0.1 M NaOH for pretreatment^70^.

### Co-culture fermentation experiments

All the stocked bacteria and yeasts isolates were grown in YPD media and then centrifuged and washed twice with fresh tea liquid (12 g/L) at 6500 g for 5 min. Synthetic, pairwise bacterium-yeast cocultures were assessed in tea liquid with 50 g/L sucrose. Bacteria species included *Komagataeibacter rhaeticus* (B_1_), *Komagataeibacter intermedius* (B_2_), *Gluconacetobacter europaeus* (B_3_), *Gluconobacter oxydans* (B_4_) and *Acetobacter senegalensis* (B_5_). Yeasts included *Brettanomyces bruxellensis* (Y_1_), *Zygosaccharomyces bailii* (Y_2_), *Candida sake* (Y_3_), *Lachancea fermentati* (Y_4_) and *Schizosaccharomyces pombe* (Y_5_). For each pairwise co-culture, the total inoculation was as a final amount at 2*10^6^ CFU/mL and the inoculation amounts of bacteria and yeast were equal. Monoculture of each species was used as control group and the inoculation was also as a final amount at 2*10^6^ CFU/mL. The cultures were then incubated at 30 °C, and microbial populations and biochemical parameters were measured after 10 d fermentation. To count bacteria and yeasts, 1000 ug/mL of natamycin or 100 mg/L chloramphenicol of was added respectively.

The B_2_-Y_1_ consortium was fermented in tea liquid supplemented with 5, 50, 100 g/L sucrose individually. To characterize the consortium, B_2_ and Y_1_ were inoculated at different initial ratios from 100:1 to 10:1, 1:1, 1:10 and 1:100. The growth rates of B_2_ and Y_1_ and the B_2_/Y_1_ ratio were calculated. Meanwhile, to determine the effect of Y_1_ on B_2_, we performed the Y_1_ monoculture experiment using the same inoculation amount as the B_2_Y_1_ co-culture. Moreover, we fixed the inoculation of B_2_ or Y_1_ (1*10^6^ CFU/mL) but varied the amount of the other species from 0 to 1*10^4^, 1*10^5^, 1*10^6^, 1*10^7^ CFU/mL. Additionally, to determine if different species differ in growth and metabolic ability, Y_1_ was co-cultured with different bacterial species (B_1_, B_2_, B_3_, B_4_ and B_5_) and B_2_ was co-cultured with different fungal species (Y_1_, Y_2_, Y_3_, Y_4_ or Y_5_) in tea substrate supplemented with 50 g/L sucrose. The population dynamics and biochemical parameters were measured at 0, 3, 6, 10 d or 0, 1, 2, 3, 6, 10 d. To count microbes in pellicles, the pellicles were first digested by shaking for 16 h at 4 °C in 15 ml of PBS buffer with 2% cellulase (Sigma Aldrich, C2730).

### Monoculture fermentation with different carbon sources

To uncover the metabolic underpinnings that drive microbial population dynamics and metabolite synthesis, we conducted a series of monoculture growth experiments for B_2_ and Y_1_ using different carbon sources. Specifically, we used 10 g/L sucrose, 10 g/L fructose, 10 g/L glucose, 50 mg/L ethanol and 2 g/L acetate for fermentation. The initial inoculation of B_2_ and Y_1_ was 2*10^6^ CFU/mL. The population and biochemical parameters were measured at 2 d intervals.

### Construction and fermentation of communities with increased complexity

The five bacterial isolates (B_1_, B_2_, B_3_, B_4_ and B_5_) and the five yeast isolates (Y_1_, Y_2_, Y_3_, Y_4_ and Y_5_) were pooled together to create a synthetic, ten-species community. In initial inoculations, all bacterial species were equally abundant, and all yeast species were also equal; however, the total bacteria-to-yeasts ratio was varied from 100:1, 10:1,1:1, 1:10, to 1:100 while fixing the total amount of inoculation (2*10^6^ CFU/mL). Two different sucrose levels, 50 and 100 g/L, were added to tea liquid for fermentation. The population dynamics and biochemical parameters were measured at 0, 3, 6, 10 d.

### Statistical analysis

All the experiments were performed for three times. Redundancy analysis between microbial community and metabolites was performed with Canoco 5.0 software (Microcomputer Power, Ithaca, NY). The hierarchical cluster analysis and principal component analysis on different consortia were performed with the SIMCA-14.1 software (Umetricus, Sweden). For hierarchical cluster analysis, the distances between observations were calculated using Ward’s method based on the concentrations of different metabolites. Heatmaps of the chemical properties of the 25 two-species fermentations and 10 single-species fermentations were produced using the heatmap package with Z-score normalization in R.

### Data availability

Amplicon sequencing data are deposited at the NCBI and available under a Bioproject ID PRJNA764354. Reference sequences of all bacterial and yeasts isolates are deposited at the NCBI. Supplementary Tables 1 and 2 contain accession numbers for all of the sequences.

## Acknowledgements

This work was supported by the National Science Foundation (1553649). X.H. was supported by the China Scholarship Council.

## Author contributions

T.L. conceived the project; T.L. and X.H. designed the study; X.H. and Y.X. performed the experiments and collected the data; X.H. and T.L. analyzed the data; T.L. and X.H. wrote the paper.

## Supplementary Information

### Supplementary Table

**Supplementary Table 1:**
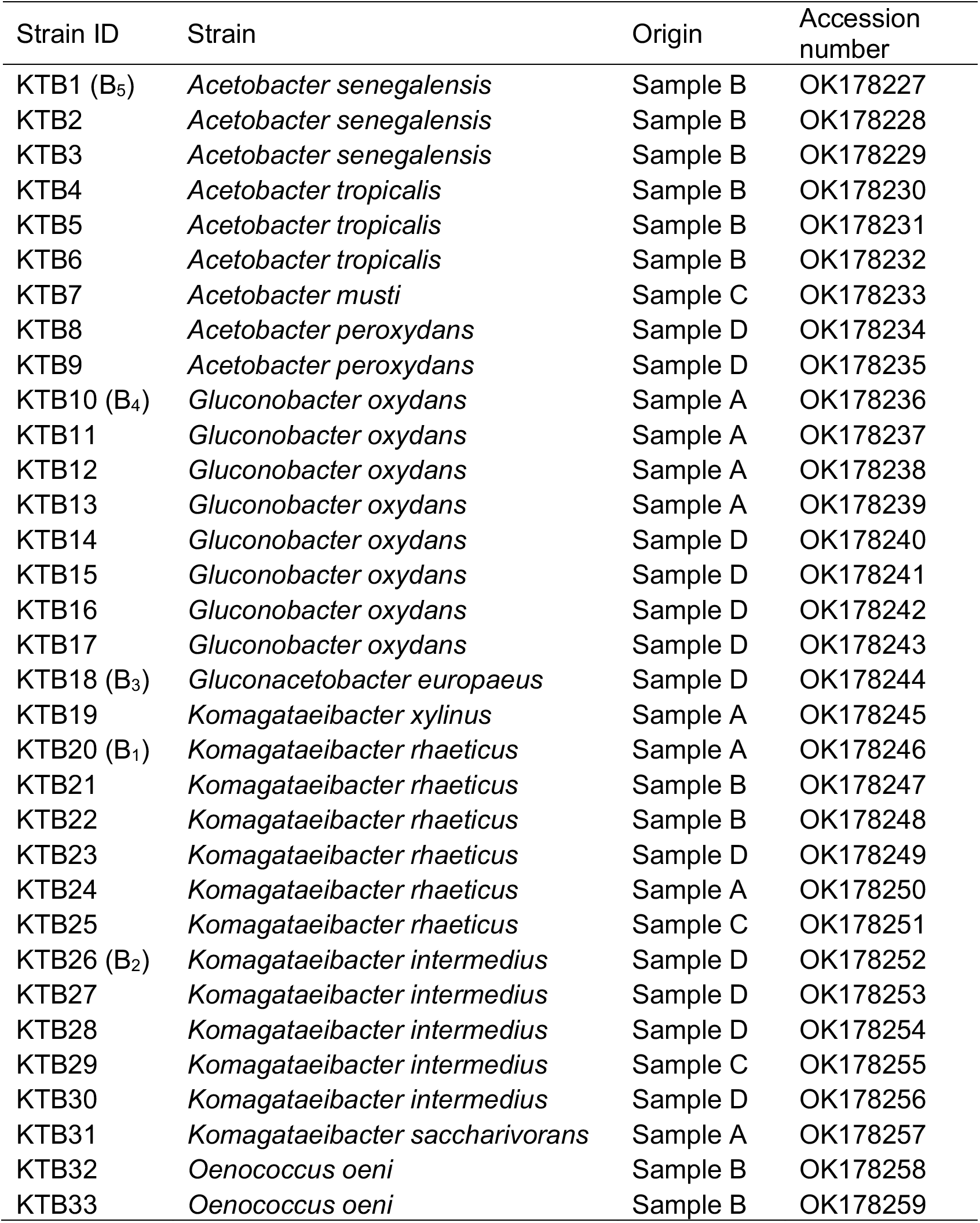
List of isolated bacterial species. Representative species diversity is given by the identification of 16S rRNA gene sequence of 33 bacterial isolates.

**Supplementary Table 2:** List of isolated fungal species. Representative species diversity is given by the identification of D1/D2 large ribosomal subunit region sequence of 30 yeast isolates.

| Strain ID | Strain | Origin | Accession number |
| --- | --- | --- | --- |
| KTY1 (Y<sub>1</sub>) | *Brettanomyces bruxellensis* | Sample A | OK271194 |
| KTY2 | *Brettanomyces bruxellensis* | Sample A | OK271195 |
| KTY3 | *Brettanomyces bruxellensis* | Sample A | OK271196 |
| KTY4 | *Brettanomyces bruxellensis* | Sample A | OK271197 |
| KTY5 | *Brettanomyces bruxellensis* | Sample D | OK271198 |
| KTY6 | *Brettanomyces bruxellensis* | Sample B | OK271199 |
| KTY7 | *Brettanomyces bruxellensis* | Sample C | OK271200 |
| KTY8 | *Brettanomyces bruxellensis* | Sample D | OK271201 |
| KTY9 | *Brettanomyces bruxellensis* | Sample A | OK271202 |
| KTY10 | *Brettanomyces bruxellensis* | Sample C | OK271203 |
| KTY11 | *Brettanomyces bruxellensis* | Sample C | OK271204 |
| KTY12 | *Brettanomyces bruxellensis* | Sample D | OK271205 |
| KTY13 | *Brettanomyces bruxellensis* | Sample D | OK271206 |
| KTY14 | *Brettanomyces bruxellensis* | Sample D | OK271207 |
| KTY15 | *Brettanomyces anomalus* | Sample D | OK271208 |
| KTY16 | *Pichia fermentans* | Sample B | OK271209 |
| KTY17 | *Pichia fermentans* | Sample B | OK271210 |
| KTY18 (Y<sub>3</sub>) | *Candida sake* | Sample B | OK271211 |
| KTY19 | *Candida sake* | Sample B | OK271212 |
| KTY20 | *Candida sake* | Sample B | OK271213 |
| KTY21 | *Candida sake* | Sample B | OK271214 |
| KTY22 | *Candida sake* | Sample B | OK271215 |
| KTY23 | *Candida sake* | Sample B | OK271216 |
| KTY24 (Y<sub>4</sub>) | *Lachancea fermentati* | Sample D | OK271217 |
| KTY25 | *Lachancea fermentati* | Sample D | OK271218 |
| KTY26 | *Lachancea fermentati* | Sample D | OK271219 |
| KTY27 | *Lachancea fermentati* | Sample D | OK271220 |
| KTY28 | *Lachancea fermentati* | Sample D | OK271221 |
| KTY29 (Y<sub>5</sub>) | *Schizosaccharomyces pombe* | Sample D | OK271222 |
| KTY30 (Y<sub>2</sub>) | *Zygosaccharomyces bailii* | Sample D | OK271223 |

**Supplementary Table 3:** Chemical property analysis of the core candidates and their controls.

|  | pH | Sucrose<br>(g/L) | Glucose<br>(g/L) | Fructose<br>(g/L) | Glucuronate<br>(g/L) | Ethanol<br>(g/L) | Acetate<br>(g/L) | Polyphenol<br>(g/L) | Flavonoid<br>(g/L) |
| --- | --- | --- | --- | --- | --- | --- | --- | --- | --- |
| B<sub>1</sub>Y<sub>1</sub> | 3.5±0.1 | 27.48±3.72 | 0.91±0.18 | 0.77±0.42 | 0 | 6.98±0.66 | 6.59±0.38 | 17.77±1.43 | 86.81±1.78 |
| B<sub>1</sub>Y<sub>2</sub> | 3.48±0.12 | 11.51±1.85 | 6.01±1 | 4.67±1.53 | 0 | 14.67±1.97 | 3.31±0.26 | 8.11±0.79 | 86.2±3.58 |
| B<sub>1</sub>Y<sub>3</sub> | 5.56±0.3 | 32.76±2.79 | 1.06±0.15 | 0 | 0 | 2.02±1.16 | 0.69±0.02 | 8.15±0.22 | 145.22±5.59 |
| B<sub>1</sub>Y<sub>4</sub> | 3.67±0.17 | 6.9±0.7 | 0 | 0.62±0.11 | 0 | 23.44±1.29 | 3.44±0.56 | 23.25±1.29 | 93.42±1.02 |
| B<sub>1</sub>Y<sub>5</sub> | 3.14±0.64 | 1.24±0.19 | 6.19±1.29 | 1.31±0.09 | 0 | 18.75±0.84 | 4.4±0.3 | 9.68±0.81 | 75.43±1.01 |
| B<sub>2</sub>Y<sub>1</sub> | 3.46±0.1 | 26.2±3.16 | 0.03±0.04 | 0.12±0.02 | 0 | 12.21±1.41 | 3.24±0.35 | 56.45±2.47 | 80.76±0.68 |
| B<sub>2</sub>Y<sub>2</sub> | 3.68±0.05 | 12.03±0.18 | 1.72±0.15 | 0.47±0.3 | 0 | 14.67±0.71 | 3.71±0.22 | 41.75±1.91 | 71.44±3.63 |
| B<sub>2</sub>Y<sub>3</sub> | 5.39±0.16 | 45.03±2.41 | 0.36±0.05 | 0.36±0.03 | 0 | 0.04±0.02 | 0 | 59.66±3.74 | 111.88±3.37 |
| B<sub>2</sub>Y<sub>4</sub> | 3.6±0.16 | 13.37±2.56 | 0.67±0.37 | 0.75±0.16 | 0 | 18.04±1.78 | 3.68±0.11 | 36.15±2.33 | 81.27±2.05 |
| B<sub>2</sub>Y<sub>5</sub> | 3.27±0.23 | 0.31±0.09 | 0.48±0.25 | 2.1±0.74 | 0 | 23.53±2.16 | 4.45±0.79 | 42.42±1.1 | 89.33±3.47 |
| B<sub>3</sub>Y<sub>1</sub> | 3.39±0.21 | 24.11±1.74 | 1.3±0.41 | 0.33±0.06 | 0 | 11.66±1.55 | 4.22±1.01 | 37.59±1.12 | 92.36±2 |
| B<sub>3</sub>Y<sub>2</sub> | 3.66±0.07 | 23.37±1.52 | 4.48±0.8 | 2.32±0.67 | 0 | 9.32±0.77 | 2.25±0.78 | 50.04±2.91 | 74.3±2.11 |
| B<sub>3</sub>Y<sub>3</sub> | 5.62±0.05 | 44.31±1.98 | 0.44±0.1 | 0 | 0 | 0.06±0.01 | 0 | 23.68±0.4 | 128.24±10.05 |
| B<sub>3</sub>Y<sub>4</sub> | 3.56±0.17 | 11.25±1.16 | 0 | 0.21±0.09 | 0 | 23.25±3.39 | 2.95±0.45 | 27.21±0.72 | 73.67±6.13 |
| B<sub>3</sub>Y<sub>5</sub> | 3.46±0.04 | 0.24±0.02 | 0 | 0.1±0.01 | 0.05±0 | 22.91±3 | 1.36±0.02 | 24.71±0.44 | 98.27±3.59 |
| B<sub>4</sub>Y<sub>1</sub> | 3.5±0.05 | 28.09±2.09 | 0.1±0.1 | 0.28±0.15 | 0.01±0 | 7.21±0.48 | 6.56±0.35 | 18.82±1.02 | 82.52±1.81 |
| B<sub>4</sub>Y<sub>2</sub> | 3.94±0.52 | 17.43±1.25 | 6.42±1.05 | 0 | 0 | 6.51±0.86 | 2.38±0.46 | 24.21±1.97 | 68.36±1.9 |
| B<sub>4</sub>Y<sub>3</sub> | 5.37±0.16 | 40.51±1.5 | 0.88±0.05 | 0.77±0.06 | 0 | 0 | 0 | 58.76±6.28 | 106.97±9.5 |
| B<sub>4</sub>Y<sub>4</sub> | 3.61±0.01 | 11.28±0.98 | 0 | 1.27±0.25 | 0.06±0.01 | 24.61±2.29 | 3.24±0.27 | 28.32±1.45 | 77.55±2.15 |
| B<sub>4</sub>Y<sub>5</sub> | 3.11±0.47 | 2.36±0.22 | 3.6±1.01 | 2.01±0.59 | 0 | 23.66±0.17 | 0.99±0.2 | 52.47±1.32 | 76.91±0.8 |
| B<sub>5</sub>Y<sub>1</sub> | 3.41±0.01 | 26.29±1.76 | 1.1±0.02 | 1.54±0.29 | 0 | 9.03±1.94 | 6.55±1.38 | 56.61±2.96 | 101.85±2.28 |
| B<sub>5</sub>Y<sub>2</sub> | 3.38±0.08 | 14.08±1.67 | 7.18±1.05 | 5±1 | 0 | 12.85±0.62 | 1.95±0.48 | 66.62±3.8 | 89.01±1.48 |
| B<sub>5</sub>Y<sub>3</sub> | 5.56±0.07 | 36.46±4.09 | 1.47±0.08 | 1.23±0.12 | 0 | 0 | 0 | 75.92±3.5 | 75.85±5.09 |
| B<sub>5</sub>Y<sub>4</sub> | 3.51±0.02 | 10.75±2.68 | 0 | 0.2±0.01 | 0 | 23.93±2.49 | 2.17±0.07 | 50.86±1.76 | 84.05±3.32 |
| B<sub>5</sub>Y<sub>5</sub> | 3.32±0.13 | 0.33±0.1 | 0 | 0.05±0.01 | 0.06±0.01 | 26.81±2.01 | 1.79±0.12 | 36.3±1.55 | 100.75±2.5 |
| B<sub>1</sub> | 4.74±0.08 | 42.57±1.21 | 0.9±0.1 | 0 | 0 | 0 | 0 | 33.22±1.81 | 122.05±3.09 |
| B<sub>2</sub> | 4.82±0.08 | 49.35±0.56 | 0 | 0 | 0 | 0 | 0 | 51.46±2.46 | 108.61±6.3 |
| B<sub>3</sub> | 4.9±0.06 | 43.47±1.06 | 0 | 0 | 0 | 0 | 0 | 47.64±3.05 | 127.14±7.31 |
| B<sub>4</sub> | 4.71±0.06 | 48.89±1.1 | 0 | 0 | 0 | 0 | 0 | 48.27±2.03 | 91.69±11.02 |
| B<sub>5</sub> | 5.27±0.05 | 43.57±1.48 | 0 | 0 | 0 | 0 | 0 | 57.29±2.28 | 116.5±15.05 |
| Y<sub>1</sub> | 3.7±0.04 | 26.46±1.03 | 0.2±0.1 | 0 | 0 | 17.54±0.58 | 0.38±0.04 | 57.2±3.53 | 110.56±1.15 |
| Y<sub>2</sub> | 3.82±0.08 | 3.62±0.65 | 0 | 0.41±0.1 | 0 | 22.21±0.58 | 0.05±0.01 | 40.95±0.71 | 56.21±8.58 |
| Y<sub>3</sub> | 5.13±0.07 | 44.11±0.21 | 0 | 0 | 0 | 0 | 0 | 42.33±1.24 | 139.23±9.59 |
| Y<sub>4</sub> | 3.77±0.08 | 1.47±0.73 | 0.45±0.05 | 0.09±0.01 | 0 | 26.37±0.88 | 0.35±0.02 | 28.03±3.21 | 101.86±4.55 |
| Y<sub>5</sub> | 3.4±0.03 | 0.19±0.06 | 0 | 1.08±0.16 | 0 | 25.73±1.02 | 0 | 34.32±1.81 | 119.05±18.16 |

### Supplementary Figures

**Supplementary Figure 1.**
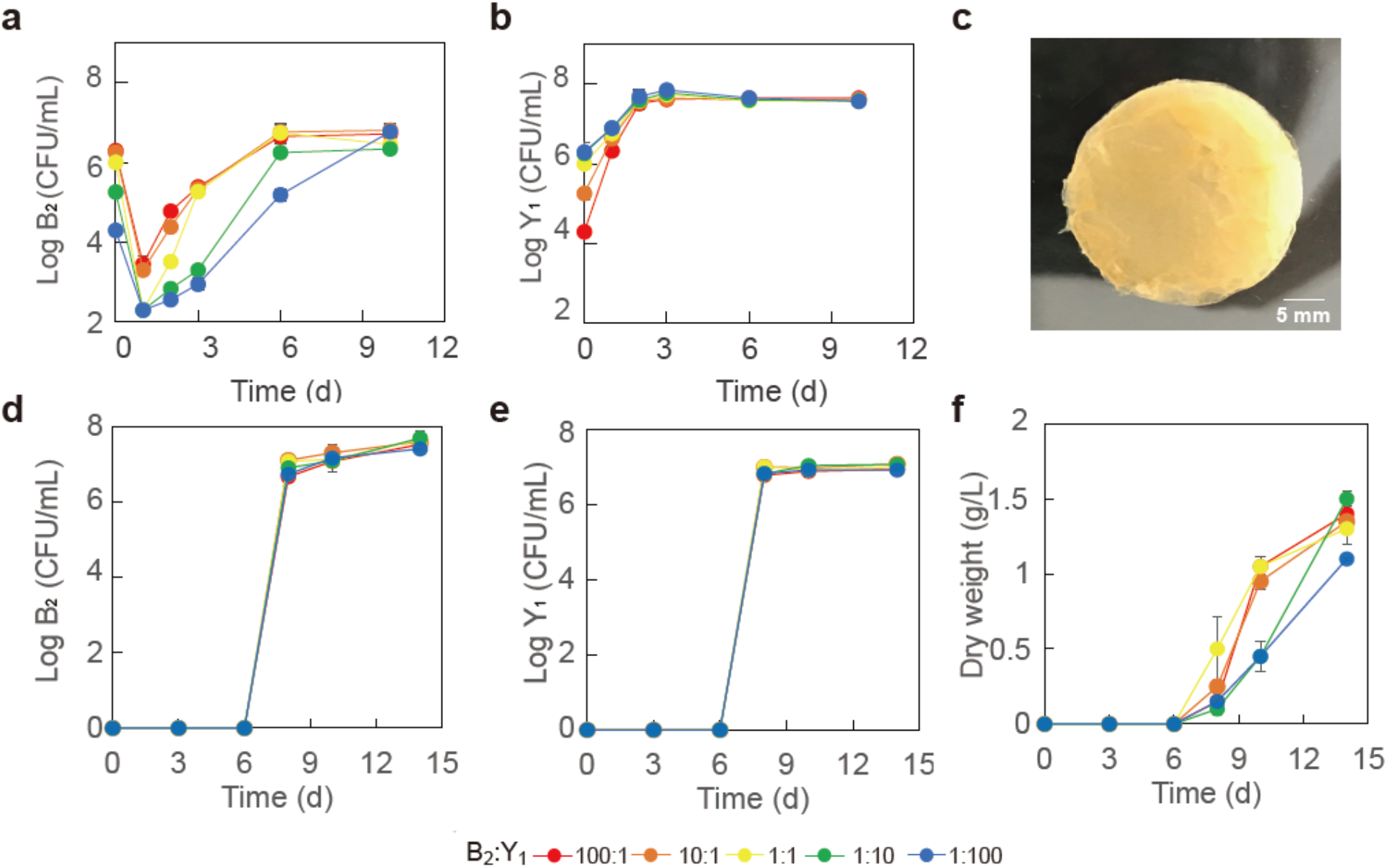
Population dynamics and pellicle formation of the minimal core (B_2_Y_1_). **a**, **b** Populations of B_2_ (**a**) and Y_1_ (**b**) in broth during a fermentation starting with 50 g/L sucrose. **c** Image of a typical pellicle formed during the fermentation. **d, e** Populations of B_2_ (**d**) and Y_1_ (**e**) in pellicle during the fermentation. **f** Dry weight of pellicles during the fermentation. Five initial compositions were used for fermentation, including 100:1 to 10:1, 1:1, 1:10 and 1:100. Bars and error bars correspond to means and s.d. respectively.

**Supplementary Figure 2.**
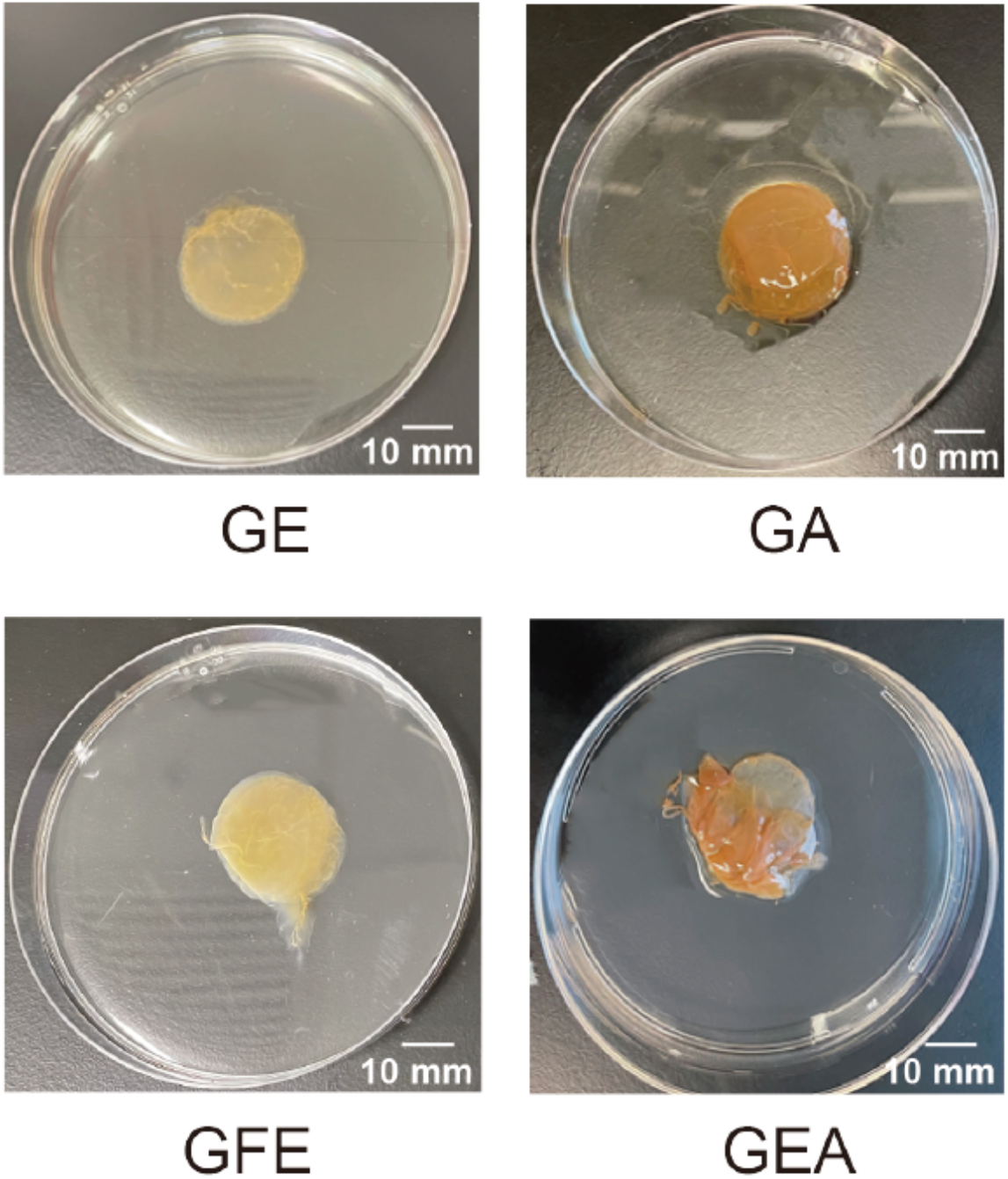
Images of pellicles formed by B_2_ monoculture with different carbon sources. GE: glucose and ethanol; GA: glucose and acetate; GFE: glucose, fructose and ethanol; GEA: glucose, ethanol and acetate.

**Supplementary Figure 3.**
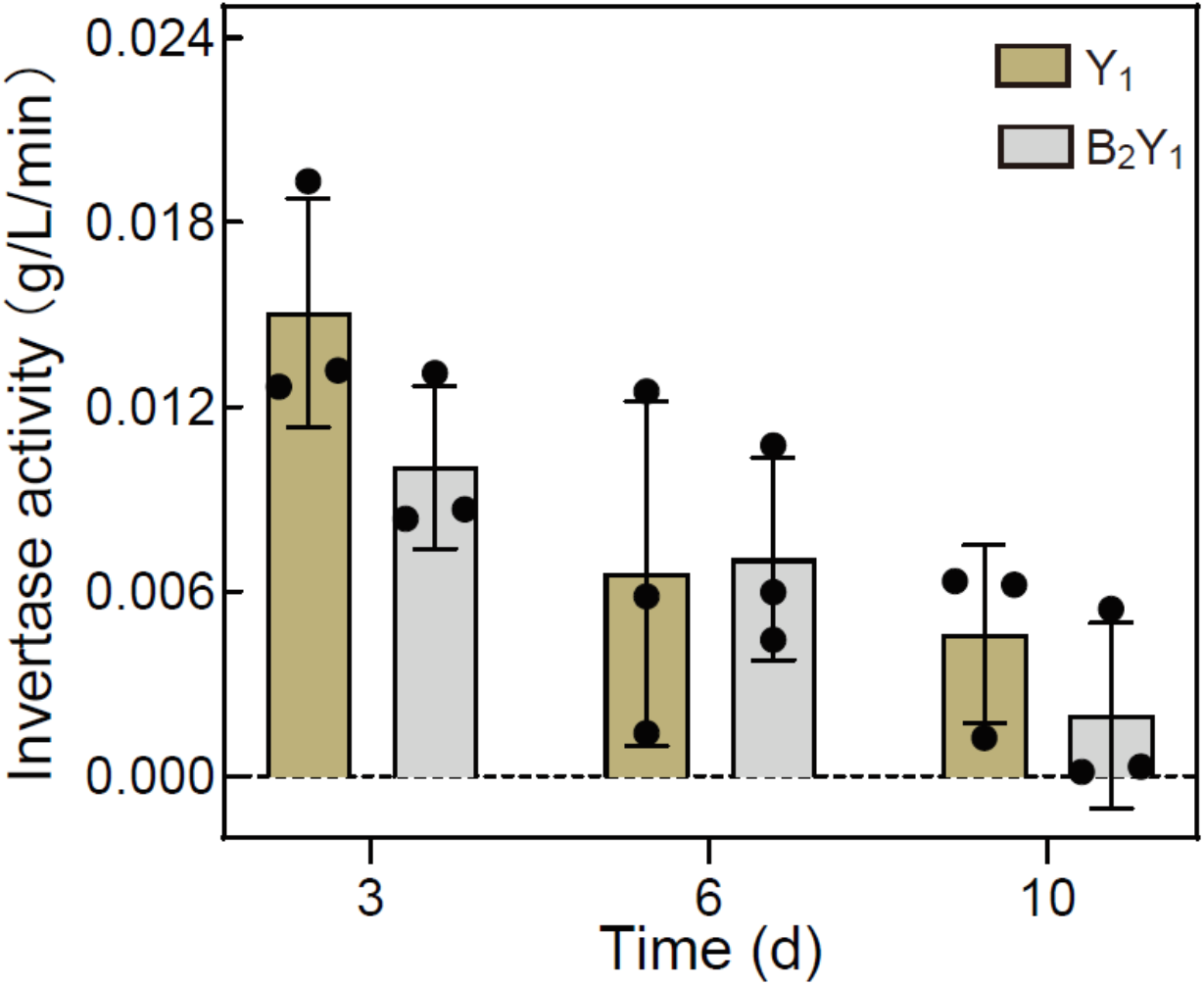
Sucrose invertase activity of Y_1_ monoculture and B_2_Y_1_ co-culture at different fermentation times. The invertase activity is defined as the amount of sucrose reduction per minute for a given amount of yeast cells (inoculation amount: 1*10^6^ CFU/mL). Bars and error bars correspond to means and s.d. respectively. T-test of paired samples in each time point did not show significant differences at *P*<0.05. (*P*=0.129732 on day 3; *P*=0.906200 on day 6; *P*=0.335084 on day 10.)

**Supplementary Figure 4.**
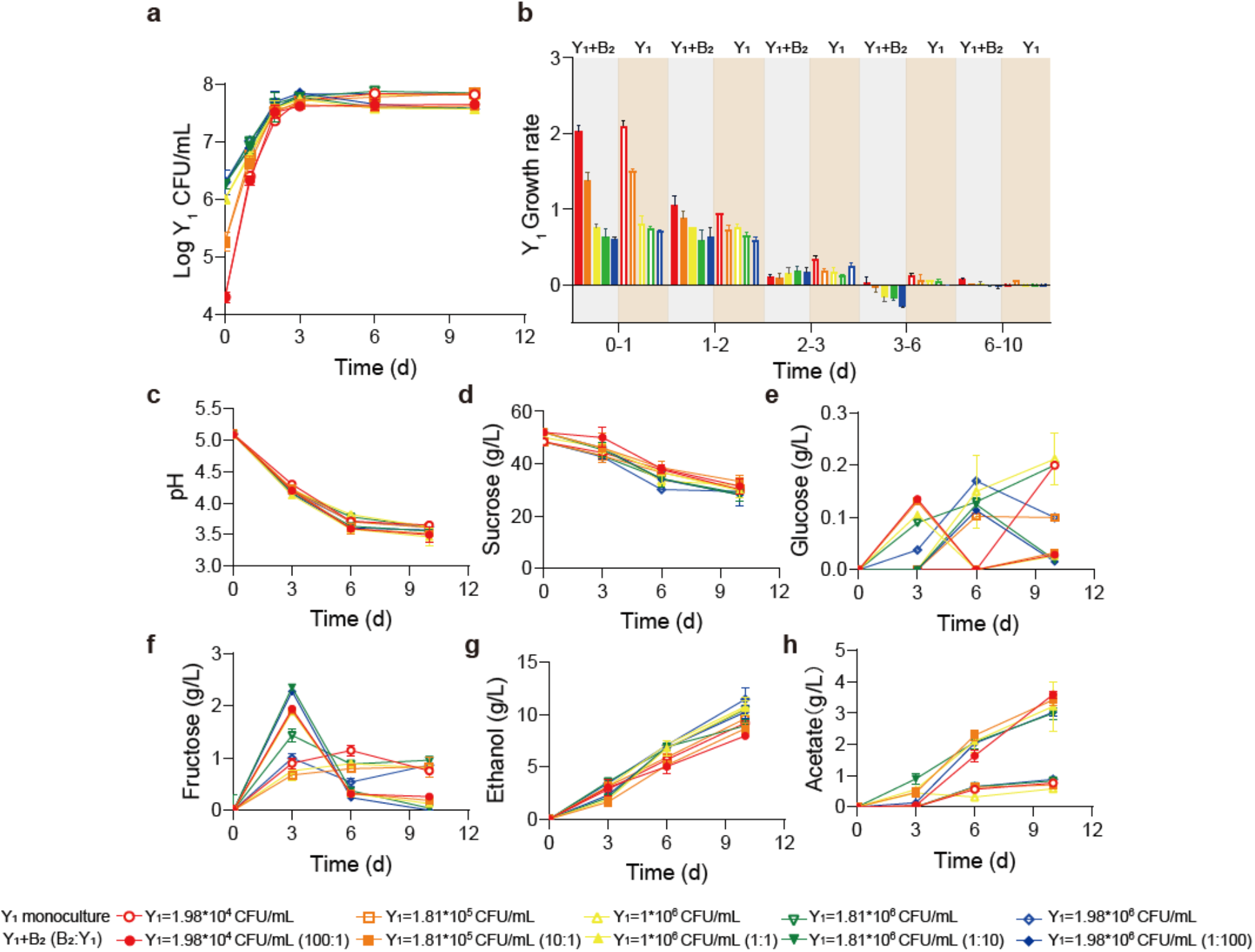
Comparation of the population and metabolic dynamics of Y_1_ monoculture and B_2_Y_1_ co-culture. **a, b** Population (**a**) and growth rate (**b**) of Y_1_ in monoculture and in the B_2_Y_1_ co-culture. **c-h** pH, carbon sources and metabolites during the fermentations of the monoculture and the co-culture. Open and filled symbols correspond to the Y_1_ monoculture and the B_2_Y_1_ co-culture, respectively. For the co-culture, the total inoculation amount was fixed at 2*10^6^ CFU/mL but the bacterium-yeast ratio was varied from 100:1 to 1:100. For the monoculture, the total bacterium population was varied in alignment with the bacterial population in the corresponding co-culture. Bars and error bars correspond to means and s.d. respectively.

**Supplementary Figure 5.**
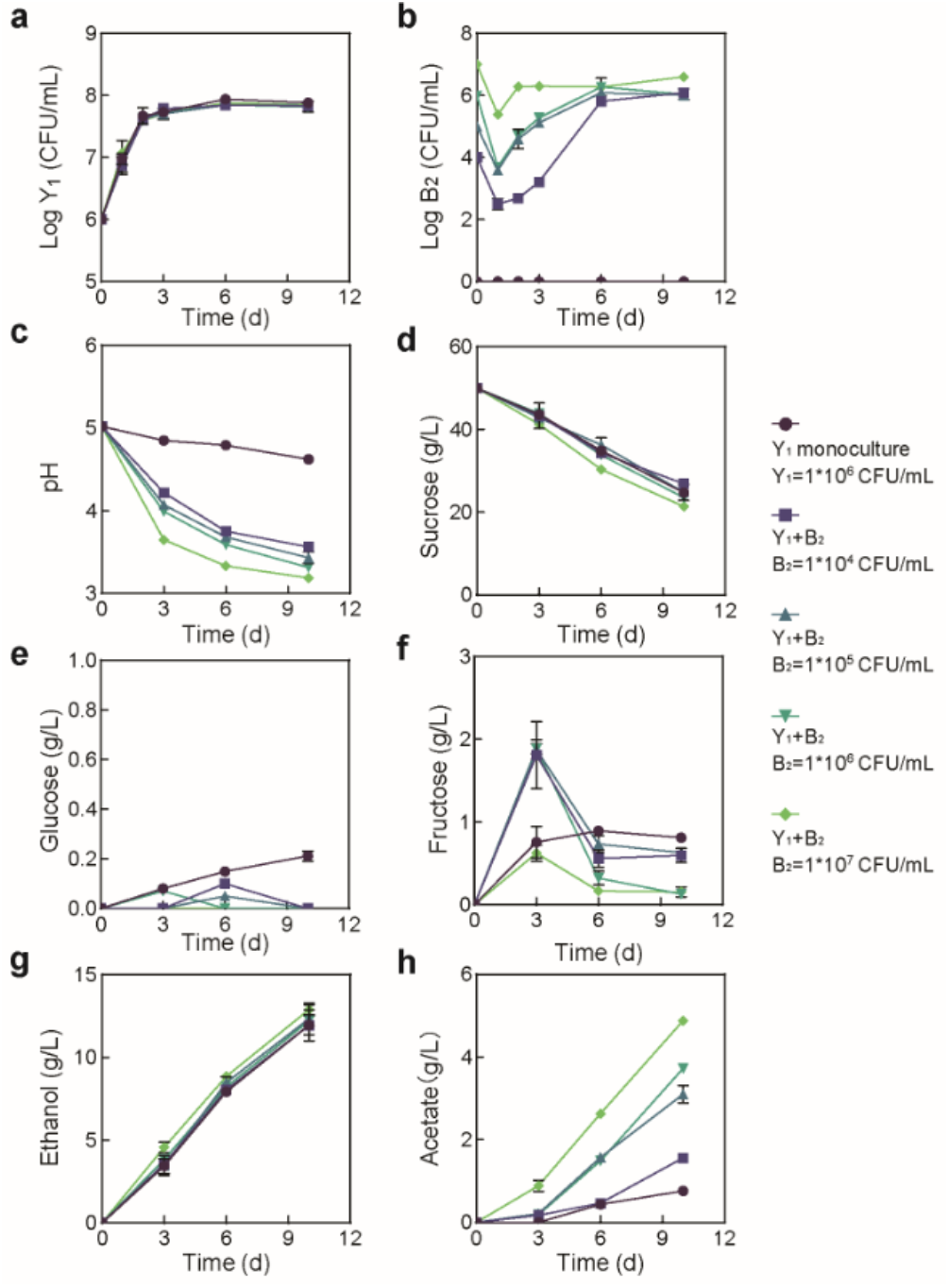
Population dynamics and metabolic profiles of the fermentations involving fixed Y_1_ and varied B_2_ initial abundances. **a** Y_1_ population dynamics. **b** B_2_ population dynamics. **c-h** pH, carbon sources and metabolites during the fermentation. The initial Y_1_ inoculation was fixed as 1*10^6^ CFU/mL but the B_2_ inoculation was varied from 0 to 1*10^4^, 1*10^5^, 1*10^6^ and 1*10^7^ CFU/mL. Bars and error bars correspond to means and s.d. respectively.

**Supplementary Figure 6.**
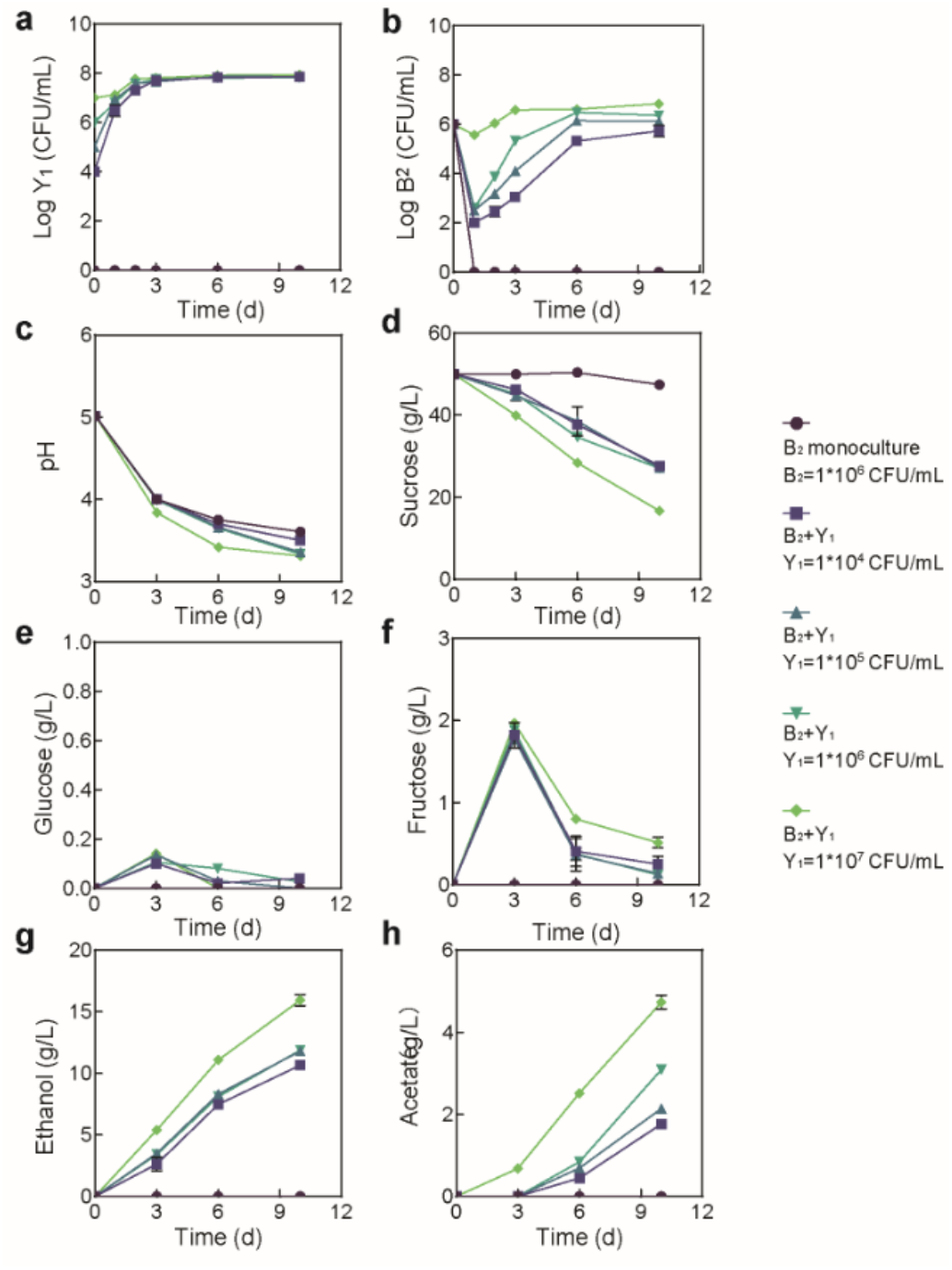
Population dynamics and metabolic profiles of the fermentations involving fixed B_2_ and varied Y_1_ initial abundances. **a** Y_1_ population dynamics. **b** B_2_ population dynamics. **c-h** pH, carbon sources and metabolites during the fermentation. The initial B_2_ inoculation was fixed as 1*10^6^ CFU/mL but the Y_1_ inoculation was varied from 0 to 1*10^4^, 1*10^5^, 1*10^6^ and 1*10^7^ CFU/mL. Bars and error bars correspond to means and s.d. respectively.

**Supplementary Figure 7.**
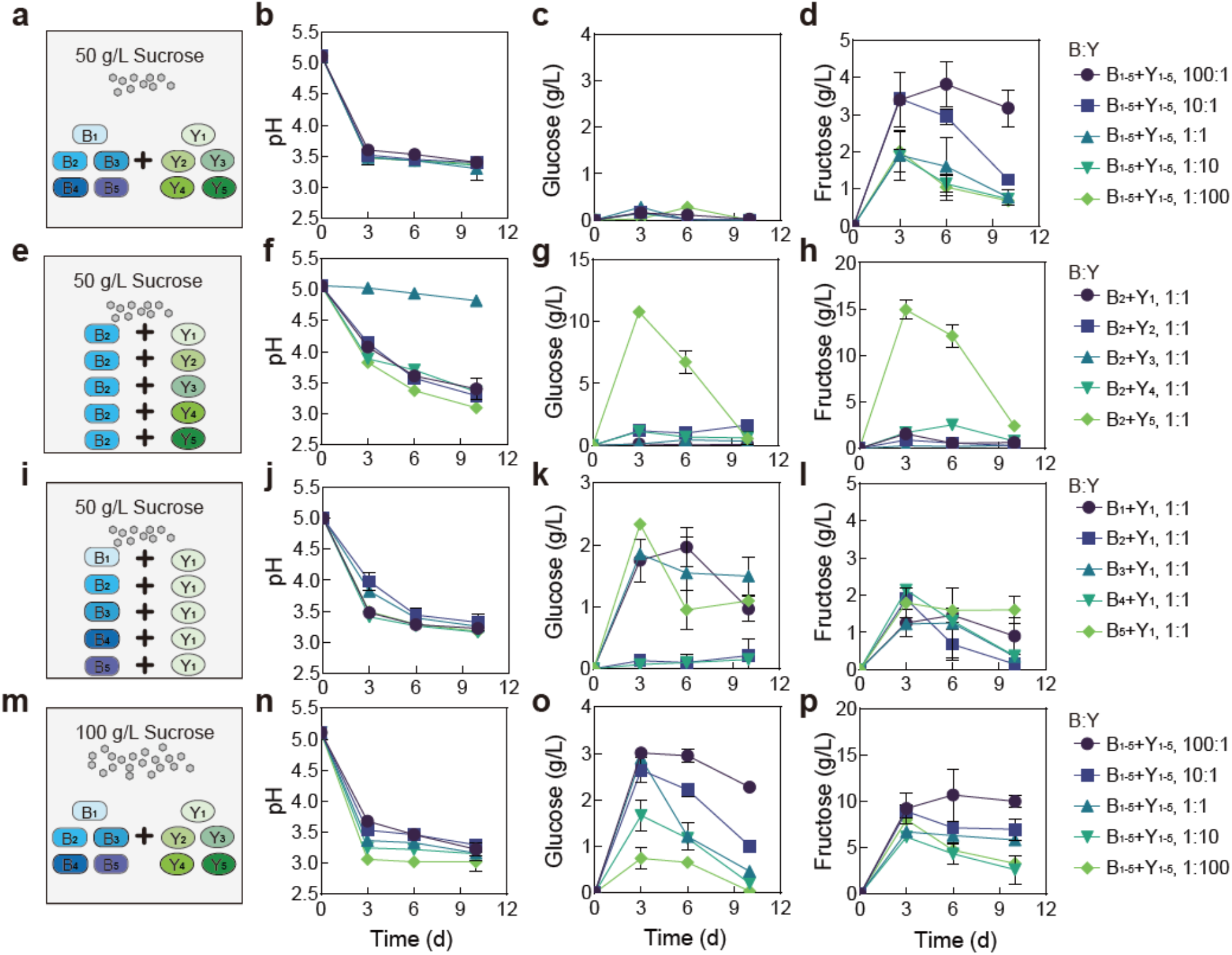
Fermentation by synthetic communities with increased complexity and altered conditions. **a** Schematic illustration of a ten-species community involving B_1_-B_5_ and Y_1_-Y_5_ in a fermentation with 50 g/L of initial sucrose. **b-d** pH (**b**), glucose (**c**) and fructose (**d**) throughout the course of the fermentation shown in **a**. **e** Schematic illustration of 5 two-species communities with each involving B_2_ and one of the yeasts (Y_1_-Y_5_) in a fermentation starting with 50 g/L sucrose. **f-h** pH, glucose and fructose during the fermentation illustrated in **e**. **I** Schematic illustration of 5 two-species communities with each involving Y_1_ and one of the bacteria (B_1_-B_5_) in a fermentation with 50 g/L of initial sucrose. **j-l** pH, glucose and fructose during the fermentation illustrated in **i**. **m** Schematic illustration of the ten-species community involving B_1_-B_5_ and Y_1_-Y_5_ in a fermentation starting with 100 g/L sucrose. **n-p** pH, glucose and fructose during the fermentation depicted in **m**. Bars and error bars correspond to means and s.d. respectively.

**Supplementary Figure 8.**
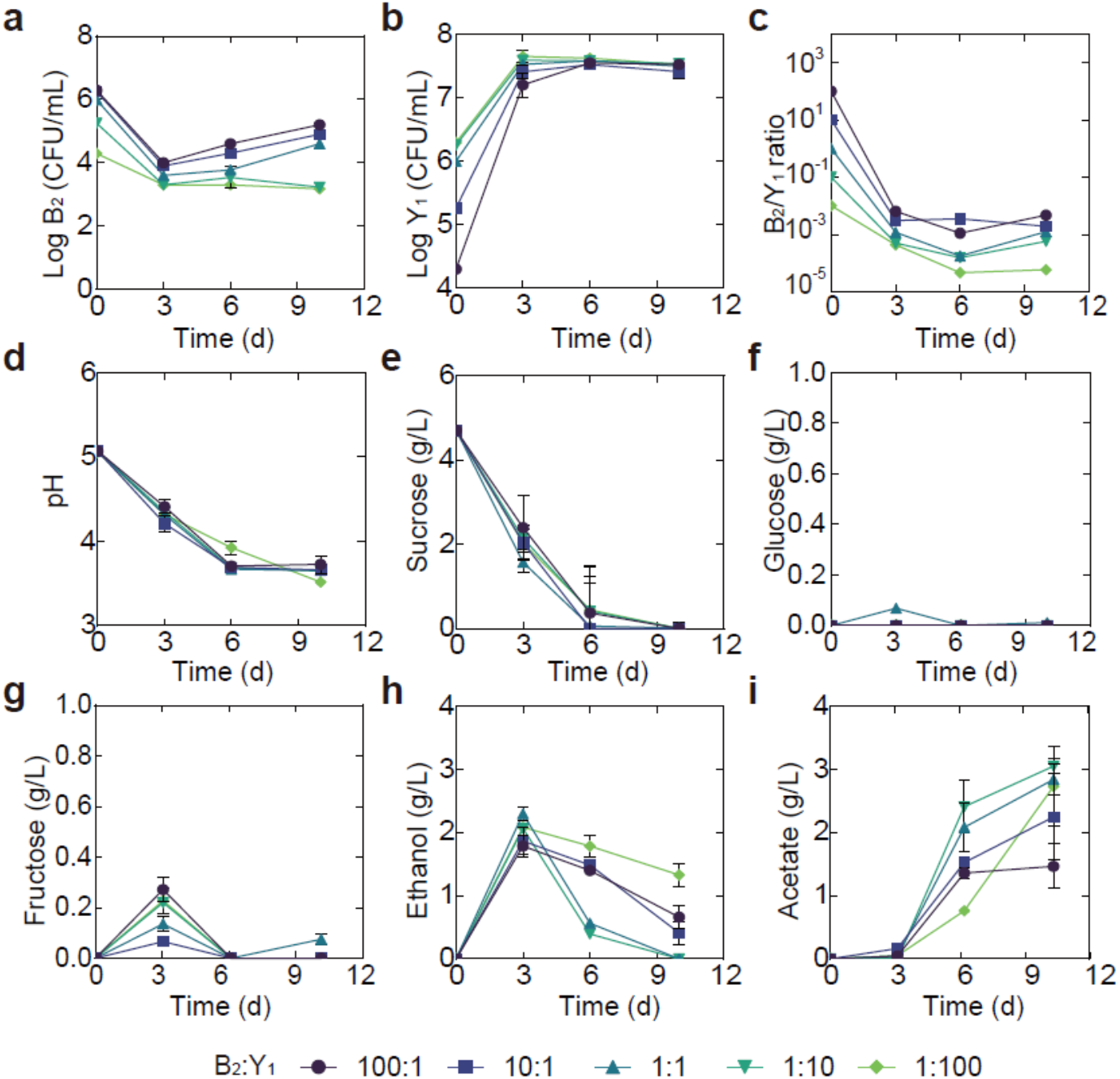
Temporal compositional and metabolic dynamics of the minimal core (B_2_Y_1_) during a fermentation with 5 g/L of initial sucrose. a, b B_2_ (a) and Y_1_ (b) population dynamics throughout the fermentation c The B_2_-to-Y_1_ ratio in the fermentation. d-i pH, carbon sources and metabolites throughout the course of the fermentation. Bars and error bars correspond to means and s.d. respectively.

**Supplementary Figure 9.**
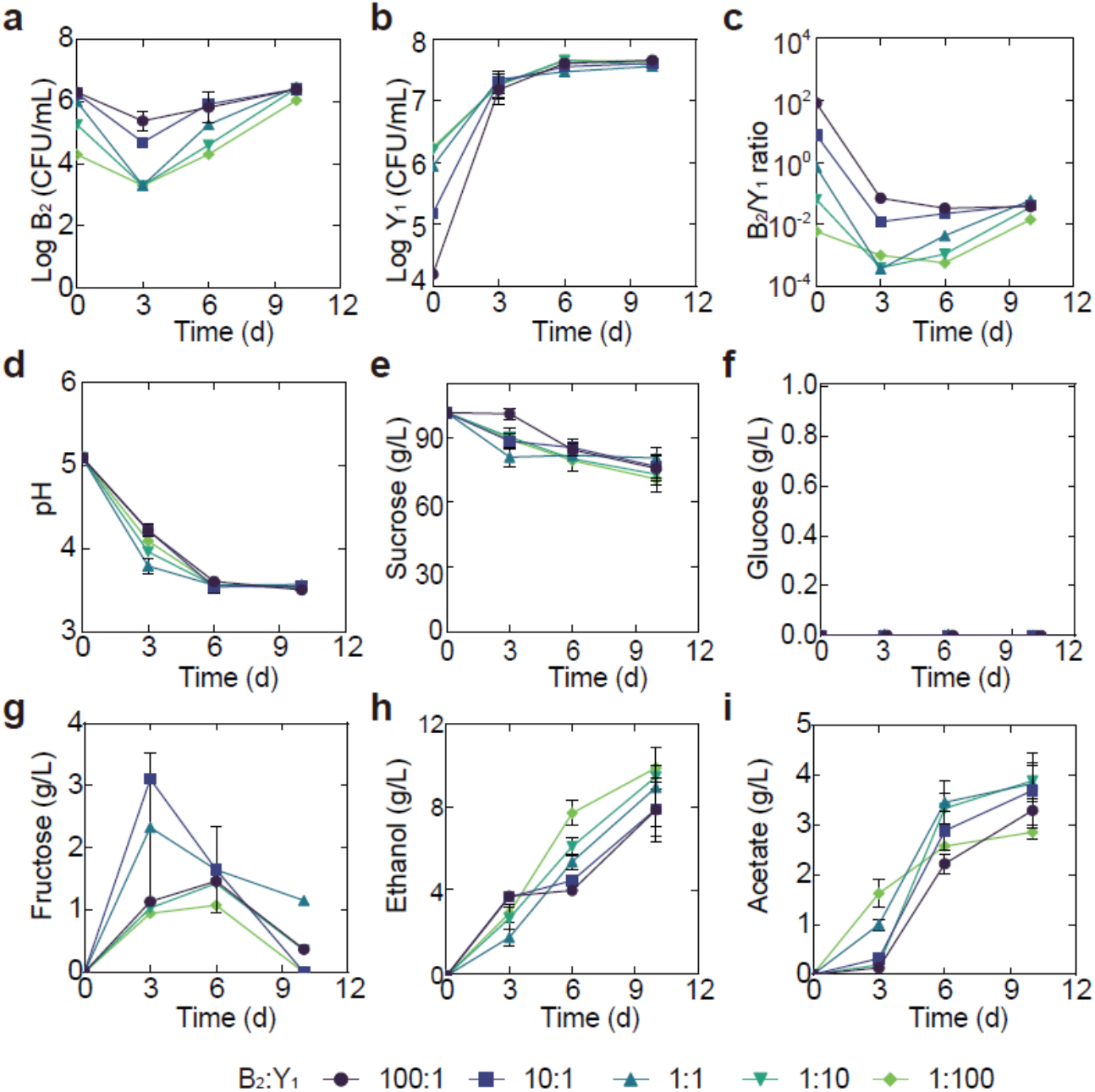
Temporal compositional and metabolic dynamics of the minimal core (B_2_Y_1_) during a fermentation with 100 g/L of initial sucrose. a, b B_2_ (a) and Y_1_ (b) population dynamics during the fermentation. c The B_2_-to-Y_1_ ratio in the fermentation. d-i pH, carbon sources and metabolites during the fermentation driven by the core. Bars and error bars correspond to means and s.d.

